## Supplementary figures and legends. for "HillTau: A fast, compact abstraction for model reduction in biochemical signaling networks"

Upinder S. Bhalla.

### Supplementary Figures

Supplementary Figure S1.

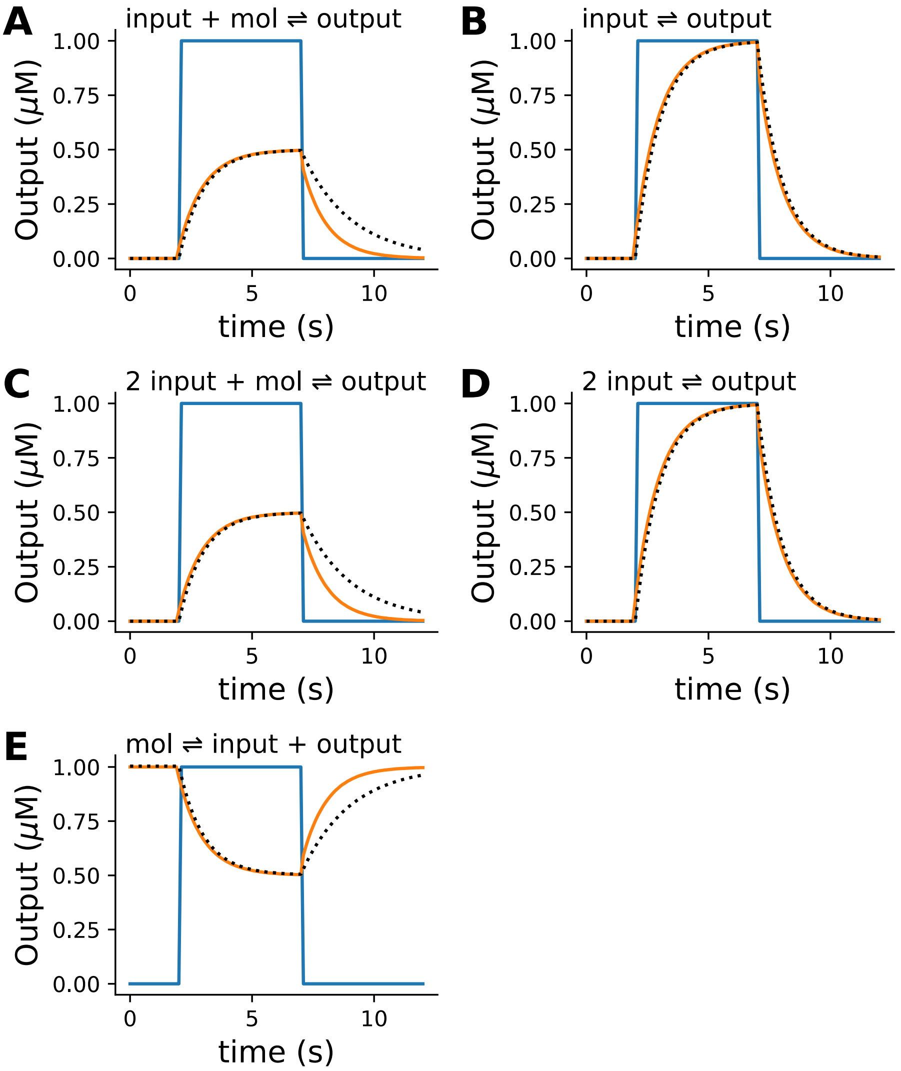


Supplementary Figure S1. HillTau fits to simple mass-action reactions indicated on top of each figure panel. Each of these is a single HillTau ‘reaction’ where ‘input’ is activator in all but Panel E, where ‘input’ is an inhibitor. In all cases the rising phase fits exactly, but in panels A, C and E the falling phase has a different time-course.

Supplementary Figure S5.1

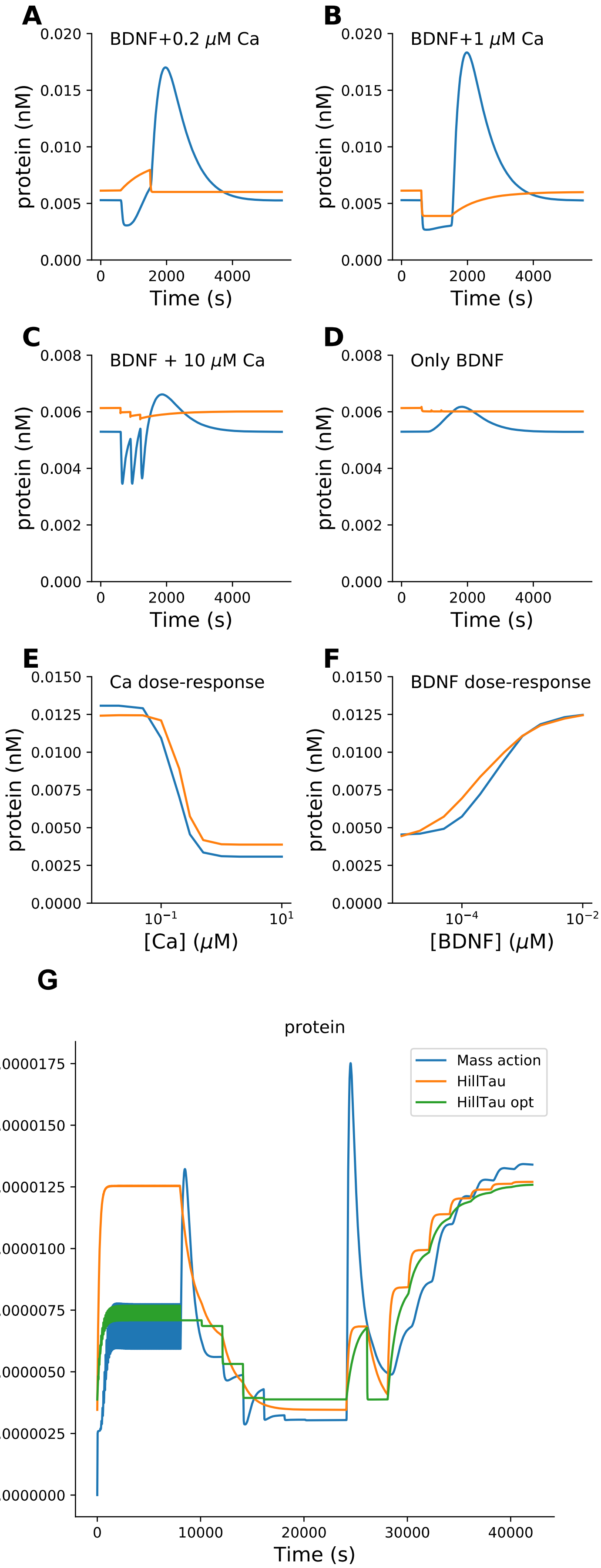


Supplementary Figure 5.1. Fit of single-reaction HillTau Model to protein synthesis pathway. Model is as in Figure 5B. Panels A-F correspond to panels D-I in Figure 5. In all cases protein production rate is readout. Blue plots are reference, orange are HillTau. A: BDNF@3.7 nM + $Ca^{2+}$@0.2 μM, 900 seconds. B: BDNF@3.7nM, $Ca^{2+}$@1μM. C: 3 pulses of BDNF@3.7 nM for 5s, coincident with $Ca^{2+}$@10μM for 1s, pulses separated by 300 s. D: Same as C, but $Ca^{2+}$ held at baseline of 0.08 μM. E: Dose-response of protein vs. $Ca^{2+}$, holding BDNF fixed at 3.7 nM. F: Dose-response of protein vs BDNF, holding $Ca^{2+}$ fixed at 0.08 μM. G: MASH optimization waveform used to fit the HillTau model for protein synthesis.

Supplementary Figure S5.2

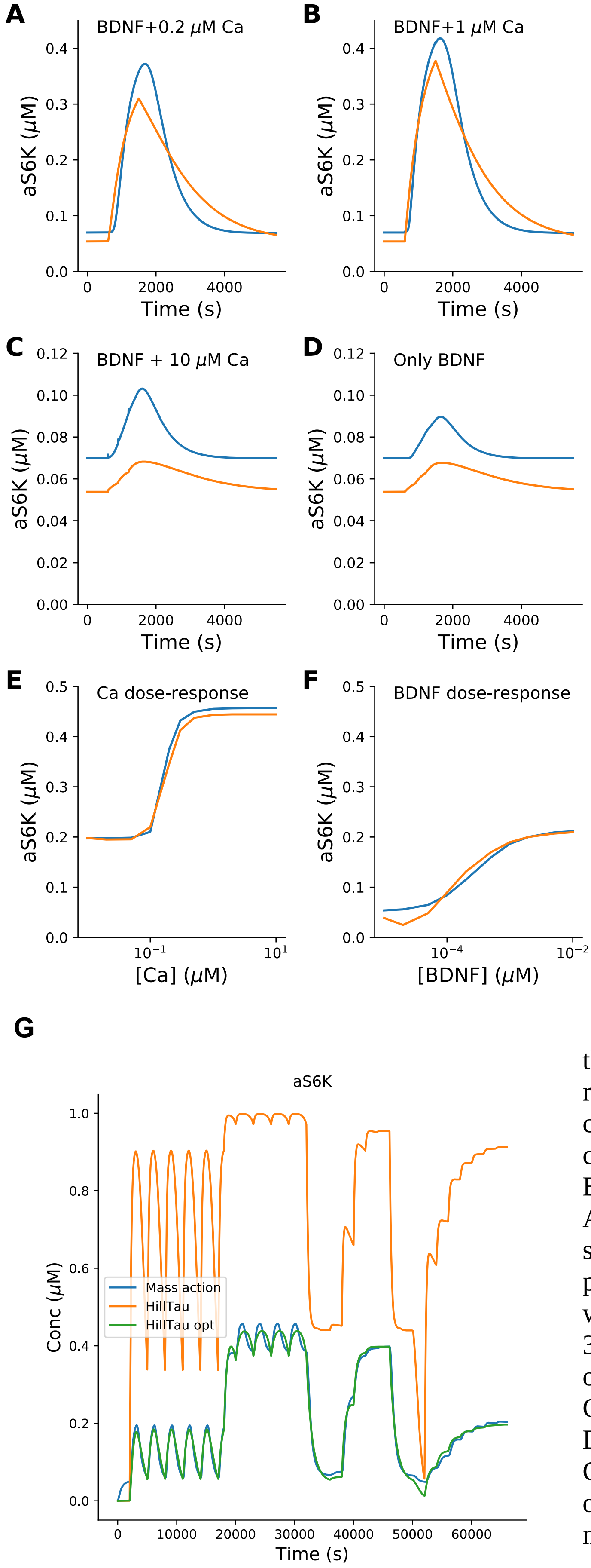


Supplementary Figure S5.2. Fitting S6K to the protein synthesis pathway model. HillTau reactions as in Figure 5C. Panels A-F correspond to panels D-I in Figure 5. In all cases activated S6K concentration is readout. Blue plots are reference, orange are HillTau. A: BDNF@3.7 nM + $Ca^{2+}$@0.2 μM, 900 seconds. B: BDNF@3.7nM, $Ca^{2+}$@1μM. C: 3 pulses of BDNF@3.7 nM for 5s, coincident with $Ca^{2+}$@10μM for 1s, pulses separated by 300 s. D: Same as C, but $Ca^{2+}$ held at baseline of 0.08 μM. E: Dose-response of protein vs. $Ca^{2+}$, holding BDNF fixed at 3.7 nM. F: Dose-response of protein vs BDNF, holding $Ca^{2+}$ fixed at 0.08 μM. G: MASH optimization waveform used to fit the HillTau model for S6K activation.

Supplementary Figure S5.3

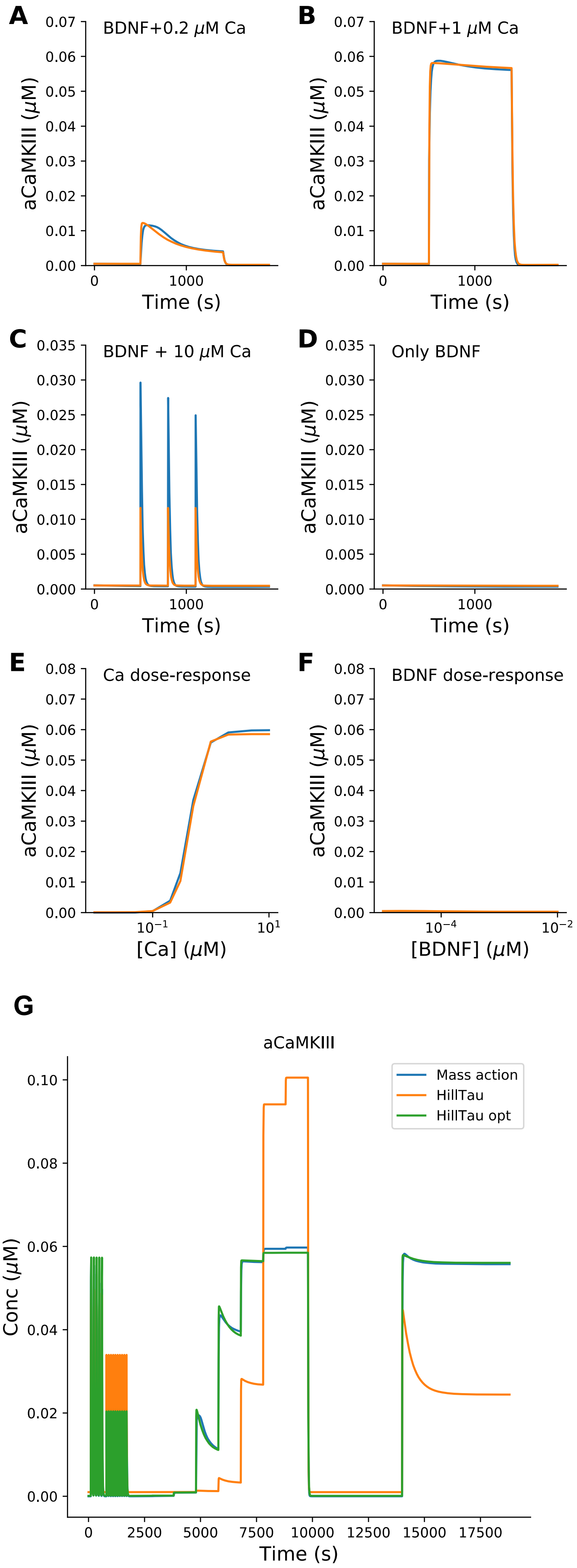


Supplementary Figure S5.3. Fitting CaMKIII to the protein synthesis pathway model. HillTau reactions as in Figure 5C. Panels A-F correspond to panels D-I in Figure 5. In all cases activated CaMKIII concentration is readout. Blue plots are reference, orange are HillTau. A: BDNF@3.7 nM + $Ca^{2+}$@0.2 μM, 900 seconds. B: BDNF@3.7nM, $Ca^{2+}$@1μM. C: 3 pulses of BDNF@3.7 nM for 5s, coincident with $Ca^{2+}$@10μM for 1s, pulses separated by 300 s. D: Same as C, but $Ca^{2+}$ held at baseline of 0.08 μM. E: Dose-response of protein vs. $Ca^{2+}$, holding BDNF fixed at 3.7 nM. F: Dose-response of protein vs BDNF, holding $Ca^{2+}$ fixed at 0.08 μM. G: MASH optimization waveform used to fit the HillTau model for CaMKIII activation.

Supplementary Figure S5.4

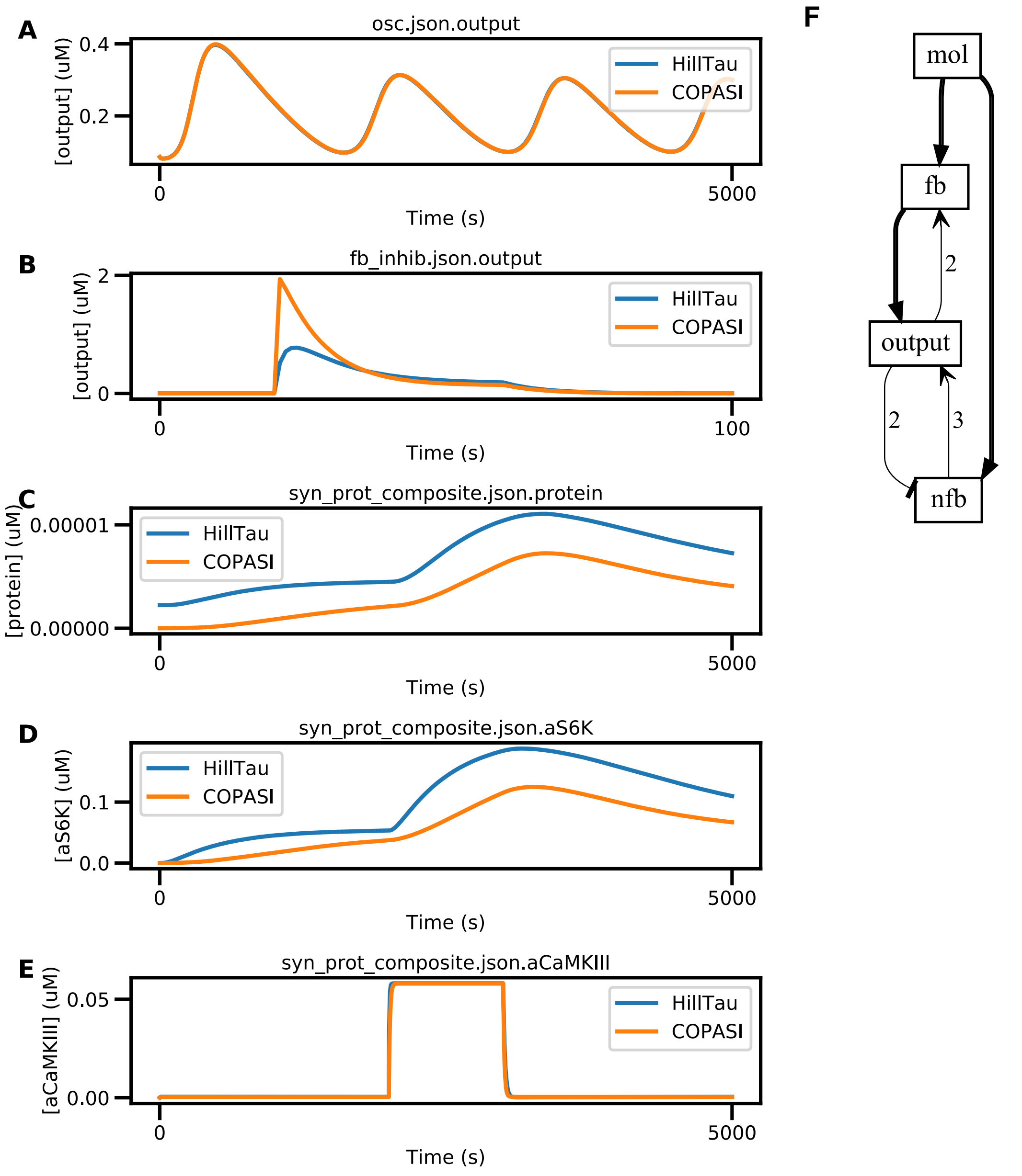


Supplementary Figure S5.4. Conversion of HillTau models to SBML, and comparison of the resultant responses simulated in HillTau and COPASI respectively. A: Oscillator model. This uses only 'tau' in its formulation, and fits to within 1%. B. Feedback inhibition model from Fig 2. A 1 uM stimulus is delivered at t = 20, and it lasts till t = 60. This has a mediocre fit or 29%. C-E: Protein synthesis model. C. Comparing protein synthesis response to a BDNF stimulus of 5 nM from t = 2000s to t = 3000s. Fit = 30% is mediocre. D. S6K activation in response to a BDNF stimulus of 5 nM from t = 2000s to t = 3000s. Fit = 26% is mediocre. E. CaMKIII activation in response to a calcium stimulus of 5 uM from t = 2000 to t = 3000s. This fits well, 1.7%. F. HillTau reaction scheme for oscillator model.

Supplementary Figure S6.1

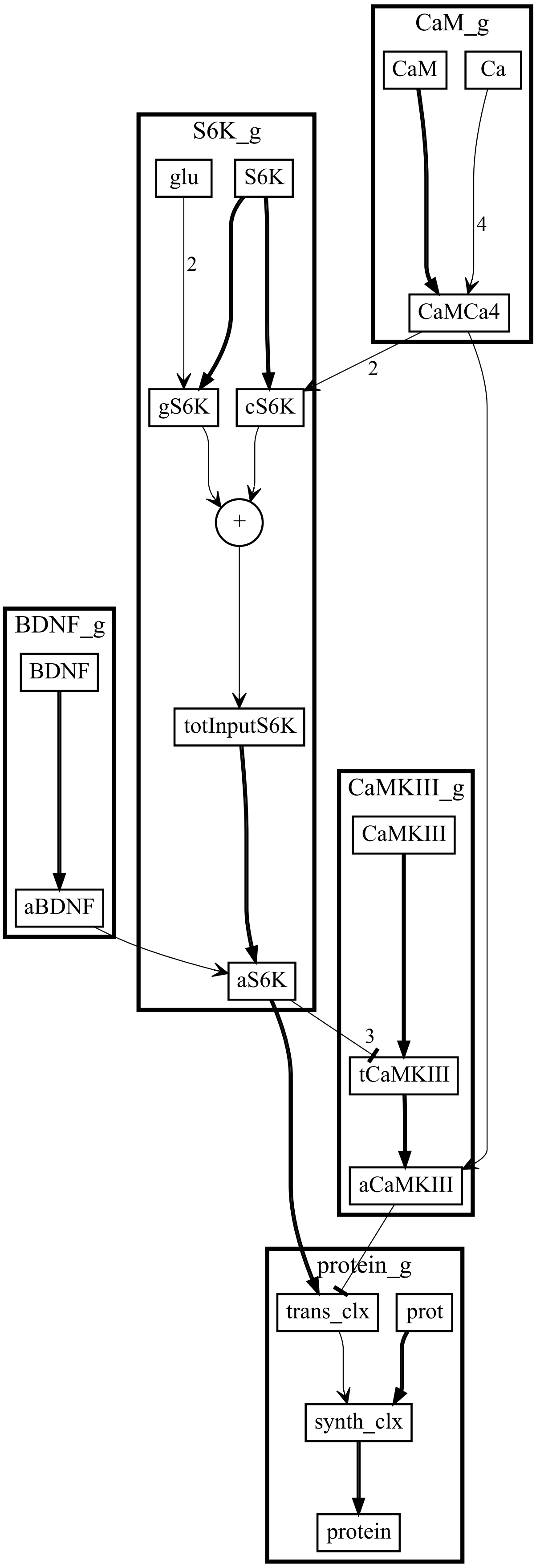


Supplementary Figure S6.1 HillTau model schematic for largest model in figure 6, with 35 HillTau parameters and 11 reactions.

Supplementary Figure S7

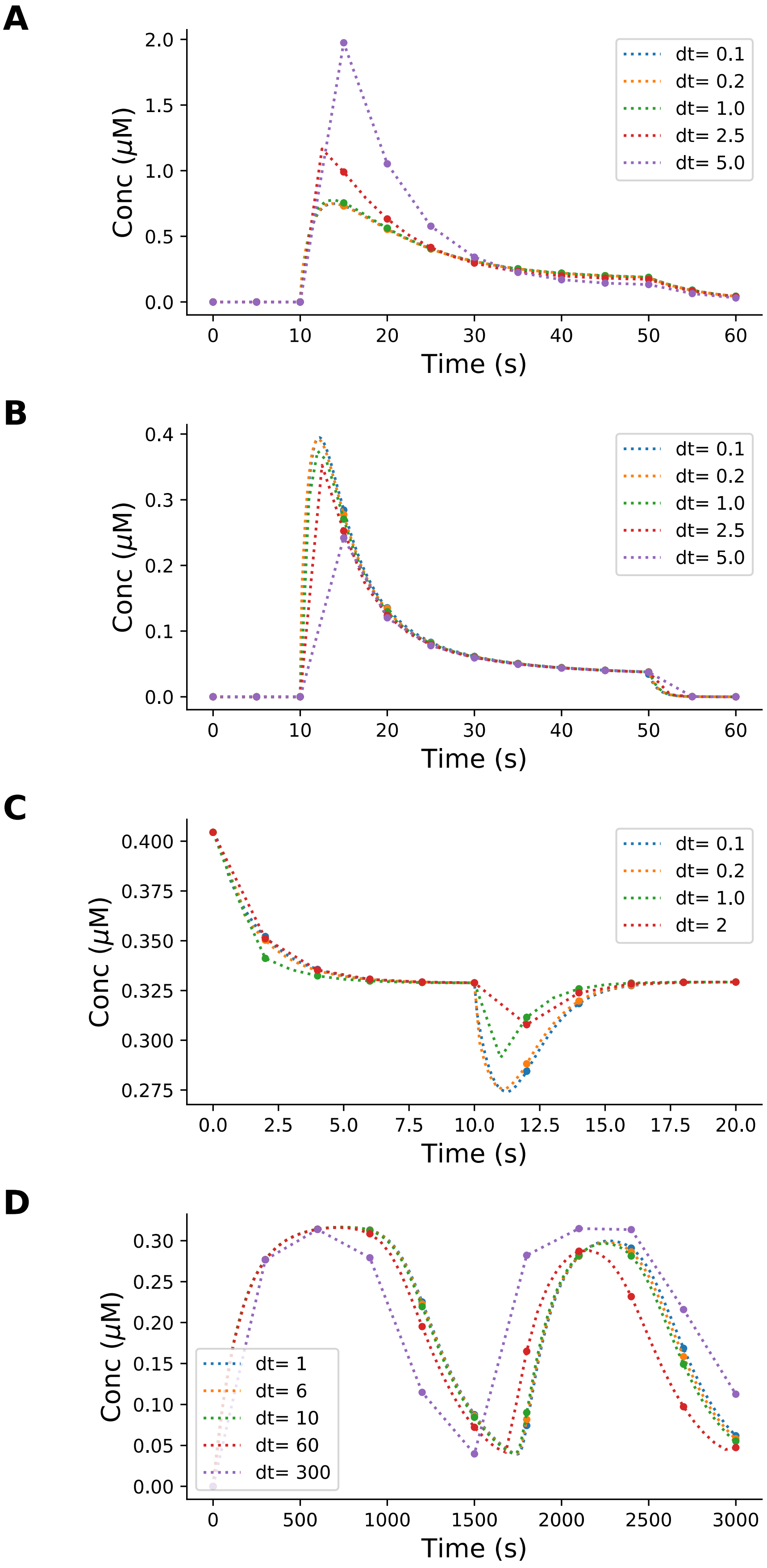

Supplementary Figure S7. Dependence of HillTau simulation output on timestep. In all panels the dashed lines represent the time-series, and the dots represent the sample points for estimating error using the smallest timestep as reference. Accuracy is reported as normalized root-mean square difference from smallest timestep. A: Feedback inhibition. Step stimulus of 1 uM is given at t = 10s, which lasts till t = 50s. 1% accuracy is achieved for dt=1s. B: feedforward inhibition. Stimulus same as A.  1.5% accuracy at dt=1s. C: BCM curve. Stimulus of 1 uM is given at t = 10s and stays till the end of the simulation. 1% accuracy at dt=1s. D: Kholodenko oscillator. Here the system is free-running. 1.2% accuracy at dt=6s.
