## Supplementary Code for "HillTau: A fast, compact abstraction for model reduction in biochemical signaling networks": conv_kkit_to_sbml.py

```
# 
# This program is free software; you can redistribute it and/or
# modify it under the terms of the GNU General Public License as
# published by the Free Software Foundation; either version 3, or
# (at your option) any later version.
# 
# This program is distributed in the hope that it will be useful,
# but WITHOUT ANY WARRANTY; without even the implied warranty of
# MERCHANTABILITY or FITNESS FOR A PARTICULAR PURPOSE.  See the GNU
# General Public License for more details.
# 
# You should have received a copy of the GNU General Public License
# along with this program; see the file COPYING.  If not, write to
# the Free Software Foundation, Inc., 51 Franklin Street, Fifth
# Floor, Boston, MA 02110-1301, USA.
# 

'''
*******************************************************************
 * File:            conv_to_sbml.py
 * Description:
 * Author:          Upinder S. Bhalla
 *
 ********************************************************************/

/**********************************************************************
** This program uses MOOSE to convert model definitions from .g to sbml.
**           copyright (C) 2020 Upinder S. Bhalla. and NCBS
**********************************************************************/
'''
from __future__ import print_function
import sys
import os
import json
import numpy as np
import argparse
import moose

def conv( fname ):
    kfile = fname + ".g"
    modelId = moose.loadModel( fname + ".g", 'model', 'none' )
    moose.writeSBML( '/model', fname + ".xml" )
    moose.delete( modelId )


def main():
    parser = argparse.ArgumentParser( description = "Convert single file or directory of files from Genesis/kkit/MOOSE .g format to SBML" )
    parser.add_argument( "filename", type = str, help= "Required: File path for source model in .g format, or for directory full of such files" )
    parser.add_argument( "-d", "--directory", action = 'store_true', help = "Optional: Flag to indicate that the program should use the path as a directory holding files to convert. If true, then the converted files will be placed in the same directory as the source files." )
    args = parser.parse_args()

    if args.directory:
        fnames = os.listdir( args.filename )
        if args.filename[-1] == '/':
            fnames = [ args.filename + i for i in fnames ]
        else:
            fnames = [ args.filename + "/" + i for i in fnames ]
    else:
        fnames = [ args.filename ]
    numConv = 0
    for f in fnames:
        if f[-2:] == ".g":
            print( "Converting '{}' to SBML.".format( f[:-2] ) )
            conv( f[:-2] )
            numConv += 1
    if numConv == 0:
        if args.directory:
            print( "Warning: no files in '{}' found with a .g suffix".format( args.filename ) )
        else:
            print( "Warning: File '{}' does not have a .g suffix".format( args.filename ) )


if __name__ == '__main__':
    main()
```
