## Supplementary Code for "HillTau: A fast, compact abstraction for model reduction in biochemical signaling networks": fig1.py

```
# 
# This program is free software; you can redistribute it and/or
# modify it under the terms of the GNU General Public License as
# published by the Free Software Foundation; either version 3, or
# (at your option) any later version.
# 
# This program is distributed in the hope that it will be useful,
# but WITHOUT ANY WARRANTY; without even the implied warranty of
# MERCHANTABILITY or FITNESS FOR A PARTICULAR PURPOSE.  See the GNU
# General Public License for more details.
# 
# You should have received a copy of the GNU General Public License
# along with this program; see the file COPYING.  If not, write to
# the Free Software Foundation, Inc., 51 Franklin Street, Fifth
# Floor, Boston, MA 02110-1301, USA.
# 

'''
*******************************************************************
 * File:            fig1.py
 * Description:
 * Author:          Upinder S. Bhalla
 *
 ********************************************************************/

/**********************************************************************
** This program demonstrates the use of HILLTAU to run elementary chemical
** reactions
**
**           copyright (C) 2020 Upinder S. Bhalla. and NCBS
**********************************************************************/
'''

import numpy as np
import matplotlib.pyplot as plt
from matplotlib import lines
import hillTau

def plotBoilerplate( panelTitle, plotPos, reacn, xlabel = '' ):
    ax = plt.subplot( 5, 2, plotPos )
    ax.spines['top'].set_visible( False )
    ax.spines['right'].set_visible( False )
    '''
    ax.tick_params( direction = 'out' )
    '''
    ax.set_xlabel( xlabel, fontsize = 14 )

    if panelTitle == "A":
        ax.set_ylabel( 'Output ($\mu$M)', fontsize = 14 )
    else:
        ax.set_ylabel( 'Conc ($\mu$M)', fontsize = 14 )
    ax.text( -0.3, 1, panelTitle, fontsize = 18, weight = 'bold', transform = ax.transAxes )
    ax.text( 0.03, 1.03, reacn, fontsize = 12, transform = ax.transAxes )
    return ax

def runSim( fname, panelTitle, reacn, plotPos ):
    jsonDict = hillTau.loadHillTau( fname )
    hillTau.scaleDict( jsonDict, hillTau.getQuantityScale( jsonDict ) )
    model = hillTau.parseModel( jsonDict )
    model.dt = 0.01

    inputMolIndex = model.molInfo.get( "input" ).index
    outputMolIndex = model.molInfo.get( "output" ).index
    
    model.advance( 2 )
    model.conc[inputMolIndex] = 1e-3
    model.advance( 2 )
    model.conc[inputMolIndex] = 0.2e-3
    model.advance( 6 )
    plotvec = np.transpose( np.array( model.plotvec ) )
    x = np.array( range( plotvec.shape[1] ) ) * model.dt
    ax = plotBoilerplate( panelTitle, plotPos, reacn, xlabel = "Time (s)" )
    ax.plot( x , 1e3*plotvec[inputMolIndex], label = "input" )
    ax.plot( x , 1e3*plotvec[outputMolIndex], label = "output" )

def plotPanelB( plotPos ):
    ax = plt.subplot( 5, 2, plotPos )
    ax.axis( 'off' )
    ax.text( -0.3, 1, "B", fontsize = 18, weight = 'bold', transform = ax.transAxes )
    ax.text( -0.1, -0.1, '...\n"Species": {\n    "input": 0.0, "mol": 1.0\n},\n"Reacs": {\n    "output": {\n        "subs": ["mol", "input"],\n        "KA": 1.0, "tau": 1.0\n    }\n}', fontsize = 12 )

def plotPanelA( plotPos ):
    KA = 1.0
    dx = 0.02
    x = np.arange( 0, 2, dx )
    y = x / (KA + x )
    ax = plotBoilerplate( "A", 1, "", xlabel = "Input ($\mu$M)" )
    ax.plot( x , y )
    ax.set_xlim( ax.get_xlim() )
    x1 = 1.0
    x2 = 0.2
    y1 = y[int(x1/dx)]
    y2 = y[int(x2/dx)]
    ax.plot( (x1,), (y1,), 'ro' )
    ax.plot( (x2,), (y2,), 'bo' )


    tau = 1.0
    x = np.arange( 0, 5, 0.1 )
    y = y1 * (1.0 - np.exp( -x / tau ) )

    ax2 = plotBoilerplate( "", 2, "", xlabel = "Time (s)" )
    ax2.set_ylabel( "" )
    ax2.set_ylim(ax.get_ylim())
    ax2.plot( x , y, "r--" )
    ax2.yaxis.set_ticklabels( [])
    x = np.arange( 0, 2, 0.02 )
    y = y1 * (1.0 - np.exp( -x / tau ) )
    ax2.plot( x , y, "g" )

    y0 = y[-1] - y2 # Find the range of the second exponential.
        # It is the level that the output has reached, to its inf value.
    x = np.arange( 0, 3, 0.02 )
    y = y0 * np.exp( -x / tau ) + y2
    ax2.plot( x + 2 , y, "b-" )

    line1 = lines.Line2D( (x1, 4), (y1, y1), color="r", linestyle = "dotted" )
    line1.set_clip_on( False )
    line2 = lines.Line2D( (x2, 4), (y2, y2), color="b", linestyle = "dotted" )
    line2.set_clip_on( False )
    ax.add_line( line1 )
    ax.add_line( line2 )
    ax2.plot( (-1, 5), (y1, y1), "r:" )
    ax2.plot( (-1, 5), (y2, y2), "b:" )


if __name__ == '__main__':
    fig = plt.figure( figsize = (7,12), facecolor='white' )
    fig.subplots_adjust( left = 0.18 )
    plotPanelA(1)
    plotPanelB(3)
    runSim( "HT_MODELS/exc.json", "C", u"input + mol \u21cc output", 4 )
    runSim( "HT_MODELS/exc2ndOrder.json", "D", u"2input + mol \u21cc output", 5 )
    runSim( "HT_MODELS/conv.json", "E", u"input \u21cc output", 6 )
    runSim( "HT_MODELS/inh.json", "F", u"mol \u21cc input + output", 7 )
    runSim( "HT_MODELS/exc_tau_baseline.json", "G", u"input+mol+0.5 \u21cc output", 8 )
    runSim( "HT_MODELS/modifier.json", "H", u"mod(input+mol) \u21cc output", 9 )
    runSim( "HT_MODELS/gain.json", "I", u"(input+mol)*gain \u21cc output", 10 )

    plt.tight_layout()
    plt.show()
```
