## Supplementary Code for "HillTau: A fast, compact abstraction for model reduction in biochemical signaling networks": fig1_supp.py

```
# 
# This program is free software; you can redistribute it and/or
# modify it under the terms of the GNU General Public License as
# published by the Free Software Foundation; either version 3, or
# (at your option) any later version.
# 
# This program is distributed in the hope that it will be useful,
# but WITHOUT ANY WARRANTY; without even the implied warranty of
# MERCHANTABILITY or FITNESS FOR A PARTICULAR PURPOSE.  See the GNU
# General Public License for more details.
# 
# You should have received a copy of the GNU General Public License
# along with this program; see the file COPYING.  If not, write to
# the Free Software Foundation, Inc., 51 Franklin Street, Fifth
# Floor, Boston, MA 02110-1301, USA.
# 

'''
*******************************************************************
 * File:            fig1_supp.py
 * Description:     Runs a series of elementary biochemical models in 
 *                  MOOSE and HillTau to illustrate how they match up.
 * Author:          Upinder S. Bhalla
 *
 ********************************************************************/
'''
from __future__ import print_function
import json
import numpy as np
import matplotlib.pyplot as plt
import moose
import hillTau

pre = 2
stim = 5
post = 5

plotDt = 0.1
stimulusAmpl = 1e-3

char = ['A', 'A', 'B', 'C', 'D', 'E', 'F', 'G', 'H' ]


def plotBoilerplate( plotPos, text ):
    ax = plt.subplot( 4, 2, plotPos )
    ax.spines['top'].set_visible( False )
    ax.spines['right'].set_visible( False )
    ax.set_xlabel( "time (s)", fontsize = 14 )

    ax.set_ylabel( 'Output ($\mu$M)', fontsize = 14 )
    ax.text( -0.35, 0.98, char[plotPos], fontsize = 18, weight = 'bold', transform = ax.transAxes )
    ax.text( 0.03, 1.01, text, fontsize = 12, transform = ax.transAxes )
    return ax

def runSim( fname, plotPos, text ):
    ht = "HT_MODELS/" + fname + ".json"
    chem = "KKIT_MODELS/" + fname + ".g"
    ax = plotBoilerplate( plotPos, text )
    modelId = moose.loadModel( chem, 'model', 'gsl' )
    #moose.le( '/model/kinetics')
    for i in range( 10, 20 ):
        moose.setClock( i, plotDt )

    iplot = moose.element( '/model/graphs/conc1/input.Co' )
    oplot = moose.element( '/model/graphs/conc1/output.Co' )
    L = moose.element( '/model/kinetics/input' )
    if moose.exists( '/model/kinetics/mol' ):
        R = moose.element( '/model/kinetics/mol' )
        R.concInit = 1e-3
    L.concInit = 0
    moose.reinit()
    moose.start( pre )
    L.concInit = stimulusAmpl
    moose.start( stim )
    L.concInit = 0
    moose.start( post )
    ivec = iplot.vector
    ovec = oplot.vector
    xvec = np.array( range( len( ivec )) ) * plotDt
    moose.delete( '/model' )

    jsonDict = hillTau.loadHillTau( ht )
    hillTau.scaleDict( jsonDict, hillTau.getQuantityScale( jsonDict ) )
    model = hillTau.parseModel( jsonDict )
    inputMolIndex = model.molInfo.get( "input" ).index
    outputMolIndex = model.molInfo.get( "output" ).index
    model.dt = plotDt
    model.reinit()
    model.advance( pre )
    model.conc[inputMolIndex] = stimulusAmpl
    model.advance( stim )
    model.conc[inputMolIndex] = 0
    model.advance( post )
    plotvec = np.transpose( np.array( model.plotvec ) )
    x = np.array( range( plotvec.shape[1] ) ) * model.dt
    ax.plot( xvec , 1000 * ivec, label = "input" )
    ax.plot( x , 1000 * plotvec[outputMolIndex], label = "output" )
    ax.plot( xvec , 1000 * ovec, "k:", label = "output" )

def main():
    fig = plt.figure( figsize = (6,9), facecolor='white' )
    fig.subplots_adjust( left = 0.18 )

    runSim( "exc", 1, u"input + mol \u21cc output" )
    runSim( "conv", 2, u"input \u21cc output" )
    runSim( "exc2ndOrder", 3, u"2 input + mol \u21cc output" )
    runSim( "conv2ndOrder", 4, u"2 input \u21cc output" )
    runSim( "inh", 5, u"mol \u21cc input + output" )
    '''
    kkitList = ["exc.g", "conv.g", "exc2ndOrder.g", "conv2ndOrder.g", "inh.g"]
    htList = ["exc.json", "conv.json", "exc2ndOrder.json", "conv2ndOrder.json", "inh.json" ]

    for j, k, h in zip( range( len( kkitList ) ), kkitList, htList ):
        runSim( k, h, j+1 )
    '''

    plt.tight_layout()
    plt.show()

if __name__ == '__main__':
    main()
```
