## Supplementary Code for "HillTau: A fast, compact abstraction for model reduction in biochemical signaling networks": fig2.py

```
# 
# This program is free software; you can redistribute it and/or
# modify it under the terms of the GNU General Public License as
# published by the Free Software Foundation; either version 3, or
# (at your option) any later version.
# 
# This program is distributed in the hope that it will be useful,
# but WITHOUT ANY WARRANTY; without even the implied warranty of
# MERCHANTABILITY or FITNESS FOR A PARTICULAR PURPOSE.  See the GNU
# General Public License for more details.
# 
# You should have received a copy of the GNU General Public License
# along with this program; see the file COPYING.  If not, write to
# the Free Software Foundation, Inc., 51 Franklin Street, Fifth
# Floor, Boston, MA 02110-1301, USA.
# 

'''
*******************************************************************
 * File:            fig2.py
 * Description:
 * Author:          Upinder S. Bhalla
 *
 ********************************************************************/

/**********************************************************************
** This program uses HILLTAU and MOOSE to compare model definitions
** in the two formalisms.
**           copyright (C) 2020 Upinder S. Bhalla. and NCBS
**********************************************************************/
'''
from __future__ import print_function
import sys
import os
import json
import re
import argparse
import numpy as np
import matplotlib.pyplot as plt
import moose
import hillTau

t1 = 20
t2 = 60
t3 = 100
i1 = 1e-3

plotDt = 1

def plotBoilerplate( panelTitle, plotPos, reacn, xlabel = 'Time (s)', ylabel = 'Conc ($\mu$M)' ):
    ax = plt.subplot( 5, 2, plotPos )
    ax.spines['top'].set_visible( False )
    ax.spines['right'].set_visible( False )
    '''
    ax.tick_params( direction = 'out' )
    '''
    ax.set_xlabel( xlabel, fontsize = 14 )
    ax.set_ylabel( ylabel, fontsize = 14 )
    ax.text( -0.32, 1, panelTitle, fontsize = 18, weight = 'bold', transform = ax.transAxes )
    #ax.text( 0.03, 1.03, reacn, fontsize = 12, transform = ax.transAxes )
    return ax

def runSim( chem, ht, plotPos ):
    modelId = moose.loadModel( chem, 'model', 'gsl' )
    stim = moose.element( '/model/kinetics/input' )
    output = moose.element( '/model/kinetics/input' )
    iplot = moose.element( '/model/graphs/conc1/input.Co' )
    oplot = moose.element( '/model/graphs/conc1/output.Co' )
    moose.setClock( iplot.tick, plotDt )
    for i in range( 10, 20 ):
        moose.setClock( i, plotDt )

    stim.concInit = 0
    moose.reinit()
    moose.start( t1 )
    stim.concInit = i1
    moose.start( t2 - t1 )
    stim.concInit = 0
    moose.start( t3 - t2 )
    ovec = oplot.vector
    x = np.array( range( len( ovec )) ) * plotDt
    ax = plotBoilerplate( "C", plotPos, "This is mass-action", xlabel = "Time (s)" )
    ax.plot( x , 1000 * ovec, label = "Mass-action" )
    moose.delete( '/model' )

    jsonDict = hillTau.loadHillTau( ht )
    hillTau.scaleDict( jsonDict, hillTau.getQuantityScale( jsonDict ) )
    model = hillTau.parseModel( jsonDict )
    model.dt = plotDt
    inputMolIndex = model.molInfo.get( "input" ).index
    outputMolIndex = model.molInfo.get( "output" ).index

    model.advance( t1 )
    model.conc[inputMolIndex] = i1
    model.advance( t2 - t1 )
    model.conc[inputMolIndex] = 0
    model.advance( t3 - t2 )
    plotvec = np.transpose( np.array( model.plotvec ) )
    x = np.array( range( plotvec.shape[1] ) ) * plotDt
    reacn = "this is ht"
    htvec = np.array( plotvec[outputMolIndex] )
    #ax = plotBoilerplate( "D", plotPos+1, reacn, xlabel = "Time (s)" )
    ax.plot( x , 1000 * htvec, label = "HillTau" )
    ax.plot( x , 1000 * plotvec[inputMolIndex], label = "input" )
    dy = htvec - ovec
    print( "Panel C normalized rms diff =", np.sqrt( np.mean( dy * dy )) / np.max( htvec ))


def runOsc( chem, ht, plotPos ):
    runtime = 6000
    modelId = moose.loadModel( chem, 'model', 'gsl' )
    #output = moose.element( '/model/kinetics/MAPK/MAPK_PP' )
    oplot = moose.element( '/model/graphs/conc1/MAPK_PP.Co' )
    moose.setClock( oplot.tick, plotDt )
    for i in range( 10, 20 ):
        moose.setClock( i, plotDt )

    moose.reinit()
    moose.start( runtime )
    ovec = oplot.vector
    x = np.array( range( len( ovec )) ) * plotDt
    ax = plotBoilerplate( "D", plotPos, "This is mass-action", xlabel = "Time (s)" )
    ax.plot( x , 1000 * ovec, label = "Mass action" )
    ax.set_ylim( 0, 0.4 )
    moose.delete( '/model' )

    jsonDict = hillTau.loadHillTau( ht )
    hillTau.scaleDict( jsonDict, hillTau.getQuantityScale( jsonDict ) )
    model = hillTau.parseModel( jsonDict )
    model.dt = plotDt
    model.reinit()
    outputMolIndex = model.molInfo.get( "output" ).index

    model.advance( runtime )
    plotvec = np.transpose( np.array( model.plotvec ) )
    x = np.array( range( plotvec.shape[1] ) ) * plotDt
    reacn = "this is ht"
    #ax = plotBoilerplate( "H", plotPos+4, reacn, xlabel = "Time (s)" )
    #ax.set_ylim( 0, 0.4 )
    htvec = np.array( plotvec[outputMolIndex] )
    ax.plot( x , 1000 * htvec, label = "HillTau" )
    dy = htvec - ovec
    print( "Panel D normalized rms diff =", np.sqrt( np.mean( dy * dy )) / np.max( htvec ))

def doseResp( model, xIndex, yIndex ):
    model.dt = plotDt
    t = 0
    x = []
    y = []
    for dose in np.exp( np.arange( -7.5, 10.0, 0.2 ) ):
        model.conc[ xIndex ] = dose
        model.advance( 1000, settle = True )
        resp = model.conc[ yIndex ]
        t += 1000
        x.append( dose )
        y.append( resp )
    return x,y

def adv( model, inputMolIndex, t, dt, val ):
    model.conc[inputMolIndex] = val
    model.advance( dt )
    t += dt
    return t

def runBis( ht, plotPos ):
    reacn = "fb vs output"
    ax = plotBoilerplate( "H", plotPos, reacn, xlabel = "fb (mM)", ylabel = "output (mM)" )
    ax.set_yscale( "log" )
    ax.set_xscale( "log" )
    ax.set_xlim( 0.0005, 10 )
    ax.set_ylim( 0.001, 5 )

    jsonDict = hillTau.loadHillTau( ht )
    reacs = jsonDict["Groups"]["output_g"]["Reacs"]
    del reacs["output"]
    hillTau.scaleDict( jsonDict, hillTau.getQuantityScale( jsonDict ) )
    model = hillTau.parseModel( jsonDict )
    outputMolIndex = model.molInfo.get( "output" ).index
    fbMolIndex = model.molInfo.get( "fb" ).index
    x, y = doseResp( model, outputMolIndex, fbMolIndex )
    ax.plot( y , x, label = "fb_vs_output", color="C6" )

    jsonDict = hillTau.loadHillTau( ht )
    reacs = jsonDict["Groups"]["output_g"]["Reacs"]
    del reacs["fb"]
    hillTau.scaleDict( jsonDict, hillTau.getQuantityScale( jsonDict ) )
    model = hillTau.parseModel( jsonDict )
    outputMolIndex = model.molInfo.get( "output" ).index
    fbMolIndex = model.molInfo.get( "fb" ).index
    x, y = doseResp( model, fbMolIndex, outputMolIndex )
    ax.plot( x , y, label = "output_vs_fb", color="C5" )

    jsonDict = hillTau.loadHillTau( ht )
    hillTau.scaleDict( jsonDict, hillTau.getQuantityScale( jsonDict ) )
    model = hillTau.parseModel( jsonDict )
    model.dt = 0.1
    fbMolIndex = model.molInfo.get( "fb" ).index
    stimMolIndex = model.molInfo.get( "stim" ).index
    outputMolIndex = model.molInfo.get( "output" ).index
    baseline = model.concInit[stimMolIndex]
    moose.reinit()
    t = 0
    t = adv( model, stimMolIndex, t, 20, baseline )
    t = adv( model, stimMolIndex, t, 1, baseline * 100 )
    t = adv( model, stimMolIndex, t, 19, baseline )
    t = adv( model, stimMolIndex, t, 5, baseline * 100 )
    t = adv( model, stimMolIndex, t, 15, baseline )
    t = adv( model, stimMolIndex, t, 1, baseline * 0.01 )
    t = adv( model, stimMolIndex, t, 19, baseline )
    t = adv( model, stimMolIndex, t, 5, baseline * 0.01 )
    t = adv( model, stimMolIndex, t, 15, baseline )
    plotvec = np.transpose( np.array( model.plotvec ) )
    x = np.array( range( plotvec.shape[1] ) ) * model.dt
    reacn = "State switching"
    ax = plotBoilerplate( "I", plotPos+1, reacn, xlabel = "Time (s)", ylabel = "output (mM)")
    ax.set_yscale( "log" )
    ax.set_ylim( 0.001, 5 )
    ax.plot( x , plotvec[fbMolIndex], label = "stimulus", color="C6" )
    ax.plot( x , plotvec[outputMolIndex], label = "output", color="C5" )

def main():
    fig = plt.figure( figsize = (6,12), facecolor='white' )
    fig.subplots_adjust( left = 0.18 )
    runSim( "KKIT_MODELS/fb_inhib.g", "HT_MODELS/fb_inhib.json", 3 )
    runOsc( "KKIT_MODELS/kholodenko.g", "HT_MODELS/kholodenko.json", 4 )
    runBis( "HT_MODELS/bistable.json", 9 )

    plt.tight_layout()
    plt.show()


if __name__ == '__main__':
    main()
```
