## Supplementary Code for "HillTau: A fast, compact abstraction for model reduction in biochemical signaling networks": fig3.py

```
# 
# This program is free software; you can redistribute it and/or
# modify it under the terms of the GNU General Public License as
# published by the Free Software Foundation; either version 3, or
# (at your option) any later version.
# 
# This program is distributed in the hope that it will be useful,
# but WITHOUT ANY WARRANTY; without even the implied warranty of
# MERCHANTABILITY or FITNESS FOR A PARTICULAR PURPOSE.  See the GNU
# General Public License for more details.
# 
# You should have received a copy of the GNU General Public License
# along with this program; see the file COPYING.  If not, write to
# the Free Software Foundation, Inc., 51 Franklin Street, Fifth
# Floor, Boston, MA 02110-1301, USA.
# 

'''
*******************************************************************
 * File:            fig3.py
 * Description:
 * Author:          Upinder S. Bhalla
 *
 ********************************************************************/

/**********************************************************************
** This program uses HILLTAU and MOOSE to compare model definitions
** specifically related to synaptic computations like the BCM curve.
**
**           copyright (C) 2020 Upinder S. Bhalla. and NCBS
**********************************************************************/
'''
from __future__ import print_function
import sys
import os
import json
import re
import argparse
import numpy as np
import matplotlib.pyplot as plt
import moose
import hillTau

plotDt = 0.1
char = ['A','A', 'B', 'C', 'D', 'E', 'F', 'G', 'H','I', 'J']

def plotBoilerplate( panelTitle, plotPos, reacn, xlabel = 'Time (s)', ylabel = 'Conc ($\mu$M)' ):
    panelX = -0.3
    ax = plt.subplot( 4, 2, plotPos )
    ax.spines['top'].set_visible( False )
    ax.spines['right'].set_visible( False )
    '''
    ax.tick_params( direction = 'out' )
    '''
    ax.set_xlabel( xlabel, fontsize = 14 )
    ax.set_ylabel( ylabel, fontsize = 14 )
    ax.text( panelX, 1.1, panelTitle, fontsize = 18, weight = 'bold', transform = ax.transAxes )
    ax.text( 0.03, 1.03, reacn, fontsize = 12, transform = ax.transAxes )
    return ax

def ts( chem, ht, ampl, plotPos, title = '' ):
    tsettle = 500 # This turns out to be needed for both models.
    tpre = 10
    tstim = 5
    tpost = 50
    modelId = moose.loadModel( chem, 'model', 'gsl' )
    Ca = moose.element( '/model/kinetics/Ca' )
    output = moose.element( '/model/kinetics/p_AMPAR' )
    iplot = moose.element( '/model/graphs/conc1/Ca.Co' )
    oplot = moose.element( '/model/graphs/conc1/p_AMPAR.Co' )
    moose.setClock( iplot.tick, plotDt )
    for i in range( 10, 20 ):
        moose.setClock( i, plotDt )

    Ca.concInit = 0.08e-3
    moose.reinit()
    moose.start( tsettle )
    moose.start( tpre )
    Ca.concInit = ampl
    moose.start( tstim )
    Ca.concInit = 0.08e-3
    moose.start( tpost )
    ivec = iplot.vector
    ovec = oplot.vector
    ivec = ivec[int( tsettle/plotDt ):]
    ovec = ovec[int( tsettle/plotDt ):]
    x = np.array( range( len( ivec )) ) * plotDt
    ax = plotBoilerplate( char[plotPos], plotPos, title, xlabel = "Time (s)", ylabel = "[synAMPAR] ($\mu$M)" )
    #ax.plot( x , 1000 * ivec, label = "input" )
    ax.plot( x , 1000 * ovec, label = "output" )
    moose.delete( '/model' )

    jsonDict = hillTau.loadHillTau( ht )
    hillTau.scaleDict( jsonDict, hillTau.getQuantityScale( jsonDict ) )
    model = hillTau.parseModel( jsonDict )
    model.dt = plotDt
    inputMolIndex = model.molInfo.get( "Ca" ).index
    outputMolIndex = model.molInfo.get( "synAMPAR" ).index

    model.advance( tpre + tsettle )
    model.conc[inputMolIndex] = ampl
    model.advance( tstim )
    model.conc[inputMolIndex] = 0.08e-3
    model.advance( tpost )
    plotvec = np.transpose( np.array( model.plotvec ) )
    x = np.array( range( plotvec.shape[1] - int(tsettle/plotDt) ) ) * plotDt
    reacn = "this is ht"
    #ax = plotBoilerplate( "B", plotPos+1, reacn, xlabel = "Time (s)" )
    #ax.plot( x , 1000 * plotvec[inputMolIndex], label = "input" )
    htvec = np.array( plotvec[outputMolIndex][int(tsettle/plotDt):] )
    ax.plot( x , 1000 * htvec, label = "output" )
    if len( htvec ) > len( ovec ):
        dy = htvec[:len(ovec)] - ovec
    else:
        dy = htvec - ovec[:len(htvec)]
    print( "time-series normalized rms diff =", np.sqrt( np.mean( dy * dy )) / np.max( htvec ))

def bisBCM( ht, plotPos ):
    ltpAmpl = 2
    ltdAmpl = 0.3
    jsonDict = hillTau.loadHillTau( ht )
    qs =  hillTau.getQuantityScale( jsonDict )
    hillTau.scaleDict( jsonDict, qs )
    model = hillTau.parseModel( jsonDict )
    model.dt = plotDt
    inputMolIndex = model.molInfo.get( "Ca" ).index
    outputMolIndex = model.molInfo.get( "synAMPAR" ).index
    model.reinit()

    origConc = model.conc[inputMolIndex]
    model.advance( 20 )
    model.conc[inputMolIndex] = ltpAmpl * qs
    model.advance( 1 )
    model.conc[inputMolIndex] = origConc
    model.advance( 29 )
    model.conc[inputMolIndex] = ltdAmpl * qs
    model.advance( 30 )
    model.conc[inputMolIndex] = origConc
    model.advance( 20 )

    plotvec = np.transpose( np.array( model.plotvec ) )
    x = np.array( range( plotvec.shape[1] ) ) * plotDt
    reacn = "ht"
    ax = plotBoilerplate( char[plotPos], plotPos, "", xlabel = "Time (s)", ylabel = "[synAMPAR] ($\mu$M)" )
    ax.plot( x , plotvec[inputMolIndex] / qs, label = "Ca", color = "C2" )
    ax.plot( x , plotvec[outputMolIndex] / qs, label = "synAMPAR", color = "C4" )
    ax.legend()

def doseResp( model, xIndex, yIndex ):
    model.dt = plotDt
    x = []
    y = []
    for dose in np.exp( np.arange( -7.0, 3.0, 0.2 ) ):
        model.conc[ xIndex ] = dose * 0.001
        model.advance( 1000, settle = True )
        resp = model.conc[ yIndex ]
        x.append( dose )
        y.append( resp * 1000 )
    return x,y

def doseRespMoose():
    x = []
    y = []
    tsettle = 200
    Ca = moose.element( '/model/kinetics/Ca' )
    output = moose.element( '/model/kinetics/p_AMPAR' )
    for i in range( 10, 20 ):
        moose.setClock( i, plotDt )
    moose.reinit()
    for dose in np.exp( np.arange( -7.0, 3.0, 0.2 ) ):
        Ca.concInit = dose * 0.001
        moose.start( tsettle )
        x.append( dose )
        y.append( output.conc * 1000 )
    moose.delete( '/model' )
    return x,y

def adv( model, inputMolIndex, t, dt, val ):
    model.conc[inputMolIndex] = val
    model.advance( dt )
    t += dt
    return t

def runDoser( kkit, ht, plotPos, title = "" ):
    ax = plotBoilerplate( char[plotPos], plotPos, title, xlabel = "[Ca] ($\mu$M)", ylabel = "[synAMPAR] ($\mu$M)" )
    ax.set_xscale( "log" )
    ax.set_xlim( 0.01, 10 )
    ax.set_ylim( 0, 0.6 )
    modelId = moose.loadModel( kkit, 'model', 'gsl' )
    x, y = doseRespMoose()
    ax.plot( x , y, label = "Syn_vs_Ca_moose" )
    moosey = np.array(y)


    jsonDict = hillTau.loadHillTau( ht )
    hillTau.scaleDict( jsonDict, hillTau.getQuantityScale( jsonDict ) )
    model = hillTau.parseModel( jsonDict )
    outputMolIndex = model.molInfo.get( "synAMPAR" ).index
    CaMolIndex = model.molInfo.get( "Ca" ).index
    x, y = doseResp( model, CaMolIndex, outputMolIndex )
    ax.plot( x , y, label = "Syn_vs_Ca" )
    hty = np.array( y )
    dy = moosey - hty
    print( "Dose-resp normalized rms diff =", np.sqrt( np.mean( dy * dy )) / np.max( hty ))


def main():
    fig = plt.figure( figsize = (6,11.1), facecolor='white' )
    fig.subplots_adjust( left = 0.18 )
    ts( "KKIT_MODELS/bcm.g", "HT_MODELS/bcm.json", 0.5e-3, 3, title = "0.5 $\mu$M Ca stim" )
    ts( "KKIT_MODELS/bcm.g", "HT_MODELS/bcm.json", 5.0e-3, 4, title = "5 $\mu$M Ca stim" )
    runDoser( "KKIT_MODELS/bcm.g", "HT_MODELS/bcm.json", 5, title = "dose-response" )
    bisBCM( "HT_MODELS/bcm_bistable.json", 7 )

    plt.tight_layout()
    plt.show()


if __name__ == '__main__':
    main()
```
