## Supplementary Code for "HillTau: A fast, compact abstraction for model reduction in biochemical signaling networks": fig5.py

```
# 
# This program is free software; you can redistribute it and/or
# modify it under the terms of the GNU General Public License as
# published by the Free Software Foundation; either version 3, or
# (at your option) any later version.
# 
# This program is distributed in the hope that it will be useful,
# but WITHOUT ANY WARRANTY; without even the implied warranty of
# MERCHANTABILITY or FITNESS FOR A PARTICULAR PURPOSE.  See the GNU
# General Public License for more details.
# 
# You should have received a copy of the GNU General Public License
# along with this program; see the file COPYING.  If not, write to
# the Free Software Foundation, Inc., 51 Franklin Street, Fifth
# Floor, Boston, MA 02110-1301, USA.
# 

'''
*******************************************************************
 * File:            fig5.py
 * Description:     Compares HillTau with ODE model for synaptic prot synth
 * Author:          Upinder S. Bhalla
 *
 ********************************************************************/

/**********************************************************************
** This program uses HILLTAU and MOOSE to compare model definitions
** in the two formalisms.
**           copyright (C) 2020 Upinder S. Bhalla. and NCBS
**********************************************************************/
'''
from __future__ import print_function
import sys
import os
import json
import re
import argparse
import numpy as np
import matplotlib.pyplot as plt
import time
import moose
import hillTau

plotDt = 1
char = ['A', 'B', 'C', 'D', 'E', 'F', 'G', 'H','I', 'J', 'K', 'L']

def plotBoilerplate( panelTitle, plotPos, reacn, xlabel = 'Time (s)', ylabel = 'Conc ($\mu$M)' ):
    panelX = -0.3
    ax = plt.subplot( 5, 2, plotPos )
    ax.spines['top'].set_visible( False )
    ax.spines['right'].set_visible( False )
    '''
    ax.tick_params( direction = 'out' )
    '''
    ax.set_xlabel( xlabel, fontsize = 14 )
    ax.set_ylabel( ylabel, fontsize = 14 )
    ax.text( panelX, 1.1, panelTitle, fontsize = 18, weight = 'bold', transform = ax.transAxes )
    ax.text( 0.03, 1.03, reacn, fontsize = 12, transform = ax.transAxes )
    return ax

def ts( chem, ht, ampl, plotPos, title = '', is_LTP = False ):
    tsettle = 6000 # This turns out to be needed for both models.
    tpre = 600 # Run for a bit to include baseline in plots.
    tpost = 4000
    Ca_rest = 0.08e-3
    BDNF_rest = 0.05e-6
    BDNF_ampl = 3.7e-6
    modelId = moose.loadModel( chem, 'model', 'gsl' )
    Ca = moose.element( '/model/kinetics/Ca' )
    BDNF = moose.element( '/model/kinetics/BDNF' )
    output = moose.element( '/model/kinetics/protein' )
    #iplot = moose.element( '/model/graphs/conc1/Ca.Co' )
    oplot = moose.element( '/model/graphs/conc1/protein.Co' )
    moose.setClock( oplot.tick, plotDt )
    for i in range( 10, 20 ):
        moose.setClock( i, plotDt )

    Ca.concInit = Ca_rest
    BDNF.concInit = BDNF_rest
    tmoose = time.time()
    moose.reinit()
    moose.start( tsettle + tpre )
    if is_LTP:
        tInter = 295
        for i in range( 3 ):
            Ca.concInit = ampl
            BDNF.concInit = BDNF_ampl
            moose.start( 1 )
            Ca.concInit = Ca_rest
            moose.start( 4 )
            BDNF.concInit = BDNF_rest
            moose.start( tInter )
        moose.start( tpost )
    else:
        tstim = 900
        Ca.concInit = ampl
        BDNF.concInit = BDNF_ampl
        moose.start( tstim )
        Ca.concInit = Ca_rest
        BDNF.concInit = BDNF_rest
        moose.start( tpost )
    tmoose = time.time() - tmoose
    ovec = oplot.vector
    ovec = ovec[int( tsettle/plotDt ):]
    x = np.array( range( len( ovec )) ) * plotDt
    ax = plotBoilerplate( char[plotPos], plotPos, title, xlabel = "Time (s)", ylabel = "protein (nM/s)" )
    ax.plot( x , 1e6 * ovec, label = "output" )
    moose.delete( '/model' )

    jsonDict = hillTau.loadHillTau( ht )
    hillTau.scaleDict( jsonDict, hillTau.getQuantityScale( jsonDict ) )
    model = hillTau.parseModel( jsonDict )
    model.dt = plotDt
    CaMolIndex = model.molInfo.get( "Ca" ).index
    BDNFMolIndex = model.molInfo.get( "BDNF" ).index
    outputMolIndex = model.molInfo.get( "protein" ).index

    tht = time.time()
    model.advance( tsettle + tpre )
    if is_LTP:
        tInter = 295
        for i in range( 3 ):
            model.conc[CaMolIndex] = ampl
            model.conc[BDNFMolIndex] = BDNF_ampl
            model.advance( 1 )
            model.conc[CaMolIndex] = Ca_rest
            model.advance( 4 )
            model.conc[BDNFMolIndex] = BDNF_rest
            model.advance( tInter )
        model.advance( tpost )
    else:
        model.conc[CaMolIndex] = ampl
        model.conc[BDNFMolIndex] = BDNF_ampl
        model.advance( tstim )
        model.conc[CaMolIndex] = Ca_rest
        model.conc[BDNFMolIndex] = BDNF_rest
        model.advance( tpost )
    tht = time.time() - tht
    plotvec = np.transpose( np.array( model.plotvec ) )
    x = np.array( range( plotvec.shape[1] - int(tsettle/plotDt) ) ) * plotDt
    reacn = "this is ht"
    htvec = np.array( plotvec[outputMolIndex][int(tsettle/plotDt):] )
    #ax = plotBoilerplate( "B", plotPos+1, reacn, xlabel = "Time (s)" )
    #ax.plot( x , 1000 * plotvec[inputMolIndex], label = "input" )
    ax.plot( x , 1e6 * htvec, label = "output" )
    #ax.plot( x , 1e6 * plotvec[outputMolIndex][int(tsettle/plotDt):], label = "output" )
    if is_LTP:
        ax.set_ylim( 0.0, 0.007 )
    else:
        ax.set_ylim( 0.0, 0.02 )
    print( "timeseries runtimes: t Moose = {:.2f};    t HillTau = {:.4f}: ".format( tmoose, tht) )
    dy = htvec - ovec
    print( char[plotPos], ": timeseries normalized rms diff =", np.sqrt( np.mean( dy * dy )) / np.max( htvec ))

def non_trained_ts( chem, ht, plotPos, title = '' ):
    tsettle = 6000 # This turns out to be needed for both models.
    tpre = 600 # Run for a bit to include baseline in plots.
    tpost = 1200
    Ca_rest = 0.08e-3
    Ca_ampl = 10e-3
    BDNF_rest = 0.05e-6
    BDNF_ampl = 3.7e-6
    modelId = moose.loadModel( chem, 'model', 'gsl' )
    Ca = moose.element( '/model/kinetics/Ca' )
    BDNF = moose.element( '/model/kinetics/BDNF' )
    output = moose.element( '/model/kinetics/protein' )
    #iplot = moose.element( '/model/graphs/conc1/Ca.Co' )
    oplot = moose.element( '/model/graphs/conc1/protein.Co' )
    moose.setClock( oplot.tick, plotDt )
    for i in range( 10, 20 ):
        moose.setClock( i, plotDt )

    Ca.concInit = Ca_rest
    BDNF.concInit = BDNF_rest
    tmoose = time.time()
    BDNF.concInit = BDNF_ampl
    moose.reinit()
    moose.start( tsettle + tpre )
    for tInter in [ 300, 120, 60, 10 ]:
        advTime = 0
        while advTime < 1200:
            Ca.concInit = Ca_ampl
            moose.start( 1 )
            Ca.concInit = Ca_rest
            moose.start( tInter - 1 )
            advTime += tInter
    moose.start( tpost )
    tmoose = time.time() - tmoose
    ovec = oplot.vector
    ovec = ovec[int( tsettle/plotDt ):]
    x = np.array( range( len( ovec )) ) * plotDt
    ax = plotBoilerplate( char[plotPos], plotPos, title, xlabel = "Time (s)", ylabel = "[protein] (nM/s)" )
    ax.plot( x , 1e6 * ovec, label = "output" )
    moose.delete( '/model' )

    jsonDict = hillTau.loadHillTau( ht )
    hillTau.scaleDict( jsonDict, hillTau.getQuantityScale( jsonDict ) )
    model = hillTau.parseModel( jsonDict )
    model.dt = plotDt
    CaMolIndex = model.molInfo.get( "Ca" ).index
    BDNFMolIndex = model.molInfo.get( "BDNF" ).index
    model.conc[BDNFMolIndex] = BDNF_ampl
    outputMolIndex = model.molInfo.get( "protein" ).index

    tht = time.time()
    model.advance( tsettle + tpre )
    for tInter in [ 300, 120, 60, 10 ]:
        advTime = 0
        while advTime < 1200:
            model.conc[CaMolIndex] = Ca_ampl
            model.advance( 1 )
            model.conc[CaMolIndex] = Ca_rest
            model.advance( tInter - 1 )
            advTime += tInter
    model.advance( tpost )
    tht = time.time() - tht
    plotvec = np.transpose( np.array( model.plotvec ) )
    x = np.array( range( plotvec.shape[1] - int(tsettle/plotDt) ) ) * plotDt
    reacn = "this is ht"
    #ax = plotBoilerplate( "B", plotPos+1, reacn, xlabel = "Time (s)" )
    #ax.plot( x , 1000 * plotvec[inputMolIndex], label = "input" )
    htvec = np.array( plotvec[outputMolIndex][int(tsettle/plotDt):] )
    #ax.plot( x , 1e6 * plotvec[outputMolIndex][int(tsettle/plotDt):], label = "output" )
    ax.plot( x , 1e6 * htvec, label = "output" )
    ax.set_ylim( 0.0, 0.02 )
    print( "non-trained timeseries runtimes: t Moose = {:.2f};    t HillTau = {:.4f}: ".format( tmoose, tht) )
    dy = htvec - ovec
    print( char[plotPos], ": timeseries normalized rms diff =", np.sqrt( np.mean( dy * dy )) / np.max( htvec ))

def doseResp( model, xIndex, yIndex, doseList ):
    model.dt = plotDt
    x = []
    y = []
    tpresettle = 5000
    model.advance( tpresettle, settle = True )
    model.conc[ xIndex ] = doseList[0] * 0.001
    for dose in doseList:
        model.conc[ xIndex ] = dose * 0.001
        model.advance( 1000, settle = True )
        resp = model.conc[ yIndex ]
        x.append( dose )
        y.append( resp * 1e6 )
    return x,y

def doseRespMoose( var, doseList ):
    x = []
    y = []
    tpresettle = 5000
    tsettle = 1000
    doseMol = moose.element( '/model/kinetics/' + var )
    output = moose.element( '/model/kinetics/protein' )
    for i in range( 10, 20 ):
        moose.setClock( i, plotDt )
    doseMol.concInit = doseList[0] * 0.001
    moose.reinit()
    moose.start( tpresettle )
    for dose in doseList:
        doseMol.concInit = dose * 0.001
        moose.start( tsettle )
        x.append( dose )
        y.append( output.conc * 1e6 )
    moose.delete( '/model' )
    return x, y

def adv( model, inputMolIndex, t, dt, val ):
    model.conc[inputMolIndex] = val
    model.advance( dt )
    t += dt
    return t

def runDoser( kkit, ht, plotPos, doseList, var = "Ca", title = "", BDNF = -1 ):
    ax = plotBoilerplate( char[plotPos], plotPos, title, xlabel = "[{}] ($\mu$M)".format( var ), ylabel = "protein (nM/s)" )
    ax.set_xscale( "log" )
    modelId = moose.loadModel( kkit, 'model', 'gsl' )
    tmoose = time.time()
    if BDNF > 0.0:
        moose.element( '/model/kinetics/BDNF' ).concInit = BDNF * 1e-3
    x, y = doseRespMoose( var, doseList )
    tmoose = time.time() - tmoose
    ax.plot( x , y, label = "prot_vs_Ca_moose" )
    mvec = np.array( y )


    jsonDict = hillTau.loadHillTau( ht )
    hillTau.scaleDict( jsonDict, hillTau.getQuantityScale( jsonDict ) )
    model = hillTau.parseModel( jsonDict )
    outputMolIndex = model.molInfo.get( "protein" ).index
    doseMolIndex = model.molInfo.get( var ).index
    if BDNF > 0.0:
        model.conc[ model.molInfo.get( "BDNF" ).index ] = BDNF * 1e-3
        model.concInit[ model.molInfo.get( "BDNF" ).index ] = BDNF * 1e-3
    tht = time.time()
    x, y = doseResp( model, doseMolIndex, outputMolIndex, doseList )
    tht = time.time() - tht
    ax.plot( x , y, label = "Prot_vs_" + var )
    ax.set_ylim( 0.0, 0.014 )
    print( "dose_resp runtimes: t Moose = {:.2f};    t HillTau = {:.4f}: ".format( tmoose, tht) )
    htvec = np.array( y )
    dy = htvec - mvec
    print( char[plotPos], ": Dose_resp normalized rms diff =", np.sqrt( np.mean( dy * dy )) / np.max( htvec ))

def main():
    fig = plt.figure( figsize = (6,12), facecolor='white' )
    fig.subplots_adjust( left = 0.18 )
    ts( "KKIT_MODELS/acc92_fixed.g", "HT_MODELS/syn_prot_composite.json", 0.2e-3, 3, title = "BDNF+0.2 $\mu$M Ca" )
    ts( "KKIT_MODELS/acc92_fixed.g", "HT_MODELS/syn_prot_composite.json", 1e-3, 4, title = "BDNF+1 $\mu$M Ca" )
    ts( "KKIT_MODELS/acc92_fixed.g", "HT_MODELS/syn_prot_composite.json", 10e-3, 5, title = "BDNF + 10 $\mu$M Ca", is_LTP = True )
    ts( "KKIT_MODELS/acc92_fixed.g", "HT_MODELS/syn_prot_composite.json", 0.08e-3, 6, title = "Only BDNF", is_LTP = True )
    CaDose = [0.01, 0.02, 0.05, 0.1, 0.2, 0.3, 0.5, 1, 2, 5, 10]
    runDoser( "KKIT_MODELS/acc92_fixed.g", "HT_MODELS/syn_prot_composite.json", 7, title = "Ca dose-response", var = "Ca", doseList = CaDose, BDNF = 3.7e-3 )
    BDNFDose = [1e-5,2e-5,5e-5,1e-4,2e-4,5e-4,1e-3,2e-3,5e-3,1e-2]
    runDoser( "KKIT_MODELS/acc92_fixed.g", "HT_MODELS/syn_prot_composite.json", 8, title = "BDNF dose-response", var = "BDNF", doseList = BDNFDose )
    non_trained_ts( "KKIT_MODELS/acc92_fixed.g", "HT_MODELS/syn_prot_composite.json", 9, title = "Pulse series" )
    runDoser( "KKIT_MODELS/acc92_fixed.g", "HT_MODELS/syn_prot_composite.json", 10, title = "Ca dose-response", var = "Ca", doseList = CaDose, BDNF = 0.05e-3 )


    plt.tight_layout()
    plt.show()


if __name__ == '__main__':
    main()
```
