## Supplementary Code for "HillTau: A fast, compact abstraction for model reduction in biochemical signaling networks": fig5_supp4.py

```
# 
# This program is free software; you can redistribute it and/or
# modify it under the terms of the GNU General Public License as
# published by the Free Software Foundation; either version 3, or
# (at your option) any later version.
# 
# This program is distributed in the hope that it will be useful,
# but WITHOUT ANY WARRANTY; without even the implied warranty of
# MERCHANTABILITY or FITNESS FOR A PARTICULAR PURPOSE.  See the GNU
# General Public License for more details.
# 
# You should have received a copy of the GNU General Public License
# along with this program; see the file COPYING.  If not, write to
# the Free Software Foundation, Inc., 51 Franklin Street, Fifth
# Floor, Boston, MA 02110-1301, USA.
# 

'''
*******************************************************************
 * File:            fig5_supp4.py
 * Description:     Converts HT models to SBML and compares outputs.
 * Author:          Upinder S. Bhalla
 *
 ********************************************************************/
'''
from __future__ import print_function
import sys
import os
import json
import re
import argparse
import numpy as np
import matplotlib.pyplot as plt
import time
sys.path.insert(1, '/home/bhalla/homework/HILLTAU/REPO/HillTau/PythonCode')
sys.path.insert(1, '/home/bhalla/homework/HILLTAU/REPO/HillTau/HT2SBML')
import hillTau
import ht2sbml
import shutil
import pycotools3
import pandas

t1 = 20
t2 = 60
t3 = 100
i1 = 1e-3
htReps = 100

plotDt = 1.0

def runHT( ht, stimMol, outputMol, events ):
    jsonDict = hillTau.loadHillTau( "HT_MODELS/"+ht )
    qs = hillTau.getQuantityScale( jsonDict )
    hillTau.scaleDict( jsonDict, qs )
    model = hillTau.parseModel( jsonDict )
    model.dt = plotDt
    htTime = 0.0
    stimIndex = model.molInfo.get( stimMol ).index
    model.reinit()
    currTime = 0.0
    for e in events:
        t0 = time.time()
        model.advance( e[0] - currTime )
        model.conc[stimIndex] = e[1] / qs
        currTime = e[0]
    htTime += time.time() - t0
    outIndex = model.molInfo.get( outputMol ).index
    return model.getConcVec( outIndex ) * 1e3

def runCOP( ht, stimMol, outputMol, events ):
    # Generate the COPASI model
    ht2sbml.conv2sbml( "HT_MODELS/"+ht, "temp.sbml", stimMol, events )

    # It croaks if you just give the file name. Have to use ./temp.sbml
    smodel = pycotools3.model.ImportSBML( "./temp.sbml" )
    m = smodel.load_model()
    #TC = pycotools3.tasks.TimeCourse( m, end = runtime, step_size = plotDt, run = True, save = False )
    TC = pycotools3.tasks.TimeCourse( m, end = events[-1][0], step_size = plotDt, run = False, report_name = 'cop.out', save = True )
    TC.run = True
    TC.set_timecourse()
    TC.set_report()
    t0 = time.time()
    sim_data = TC.simulate()
    ctime = time.time() - t0
    ret = pandas.read_csv( 'cop.out', sep = '\t' )
    ret.style.set_properties(**{ 'font-size': '20pt', })
    return np.array( ret["["+outputMol+"]"] ) * 1e6

def runBoth( plotPos, ht, stimMol, outputMol, events ):
    for i in range( len( events ) ):
        events[i][1] *= 1e-3
    hvec = runHT( ht, stimMol, outputMol, events )
    cvec = runCOP( ht, stimMol, outputMol, events )
    print( "HVec shape = ", hvec.shape, "       CVEC shape = ", cvec.shape )
    plt.rcParams.update({'font.size': 14})
    ax = plt.subplot( 5, 1, plotPos )
    ax.plot( hvec, label = "HillTau" )
    ax.plot( cvec, label = "COPASI" )
    panel = ["A", "A", "B", "C", "D", "E"]
    ax.text( -0.18, 1, panel[plotPos], fontsize = 18, weight = 'bold', transform = ax.transAxes )
    plt.xticks( fontsize = 14 )
    plt.yticks( fontsize = 14 )
    ax.set_xlabel( "Time (s)", fontsize = 14 )
    ax.set_ylabel( "[{}] (uM)".format( outputMol ), fontsize = 14 )
    ax.set_title( ht + "." + outputMol, fontsize = 14 )
    ax.legend( fontsize = 14 )
    dy = hvec - cvec
    print( "Panel {} normalized rms diff = {}".format( panel[plotPos], np.sqrt( np.mean( dy * dy )) / np.max( hvec )) )


def main():
    fig = plt.figure( figsize = (8, 12) )
    runBoth( 1, "osc.json", "output", "output", [[5000, 0]] )
    runBoth( 2, "fb_inhib.json", "input", "output", [[20,1], [60,0], [100,0]] )
    runBoth( 3, "syn_prot_composite.json", "BDNF", "protein", [[2000,5e-6], [3000, 50e-9], [5000, 50e-9]] )
    runBoth( 4, "syn_prot_composite.json", "BDNF", "aS6K", [[2000,5e-6], [3000, 50e-9], [5000, 50e-9]] )
    runBoth( 5, "syn_prot_composite.json", "Ca", "aCaMKIII", [[2000,5e-3], [3000,0.08e-3], [5000,0.08e-3]] )
    plt.tight_layout()
    plt.show()

if __name__ == '__main__':
    main()
```
