## Supplementary Code for "HillTau: A fast, compact abstraction for model reduction in biochemical signaling networks": fig6.py

```
# 
# This program is free software; you can redistribute it and/or
# modify it under the terms of the GNU General Public License as
# published by the Free Software Foundation; either version 3, or
# (at your option) any later version.
# 
# This program is distributed in the hope that it will be useful,
# but WITHOUT ANY WARRANTY; without even the implied warranty of
# MERCHANTABILITY or FITNESS FOR A PARTICULAR PURPOSE.  See the GNU
# General Public License for more details.
# 
# You should have received a copy of the GNU General Public License
# along with this program; see the file COPYING.  If not, write to
# the Free Software Foundation, Inc., 51 Franklin Street, Fifth
# Floor, Boston, MA 02110-1301, USA.
# 

'''
*******************************************************************
 * File:            fig6.py
 * Description:     Runs a series of models in MOOSE, COPASI and HillTau to 
 *                  compare their runtimes.
 * Author:          Upinder S. Bhalla
 *
 ********************************************************************/
'''
from __future__ import print_function
import sys
import os
import json
import re
import argparse
import numpy as np
import matplotlib.pyplot as plt
import time
import moose
import hillTau
import shutil
import pycotools3

t1 = 20
t2 = 60
t3 = 100
i1 = 1e-3
htReps = 100

plotDt = 1.0

modelList = [
    ["exc.g", "exc.json", "exc.xml", 1e5],
    ["conv.g", "conv.json", "conv.xml", 1e5],
    ["fb_inhib.g", "fb_inhib.json", "fb_inhib.xml", 1e5],
    ["kholodenko.g", "kholodenko.json", "kholodenko.xml", 1e5],
    ["bcm.g", "bcm.json", "bcm.xml", 1e5],
    ["acc92_fixed.g", "syn_prot_composite.json", "acc92_fixed.xml", 1e4],
    ["autsim_v2_17Jul2020.g", "aut6.json", "autsim_v1_17Jul2020.xml", 1e3],
]

def runSim( chem, ht, cps, runtime ):
    print( "-------------------------------------> RUNNING: ", chem )
    modelId = moose.loadModel( "KKIT_MODELS/" + chem, 'model', 'gsl' )
    for i in range( 0, 20 ):
        moose.setClock( i, plotDt )

    moose.reinit()
    mooseTime = time.time()
    moose.start( runtime )
    mooseTime = time.time() - mooseTime
    moose.delete( '/model' )

    jsonDict = hillTau.loadHillTau( "HT_MODELS/" + ht )
    hillTau.scaleDict( jsonDict, hillTau.getQuantityScale( jsonDict ) )
    model = hillTau.parseModel( jsonDict )
    model.dt = plotDt
    htTime = 0.0
    for i in range( htReps ):
        model.reinit()
        t0 = time.time()
        model.advance( runtime )
        htTime += time.time() - t0

    # Now run it again, but in steady-state mode for HillTau
    htTime2 = 0.0
    for i in range( htReps ):
        model.reinit()
        t0 = time.time()
        model.advance( runtime, settle = True )
        htTime2 += time.time() - t0

    # Now do the COPASI thing.
    shutil.copy( "SBML_MODELS/" + cps, "temp.sbml" )

    # It croaks if you just give the file name. Have to use ./temp.sbml
    smodel = pycotools3.model.ImportSBML( "./temp.sbml" )
    m = smodel.load_model()
    #TC = pycotools3.tasks.TimeCourse( m, end = runtime, step_size = plotDt, run = True, save = False )
    TC = pycotools3.tasks.TimeCourse( m, end = runtime, step_size = plotDt, run = False, save = False )
    TC.run = True
    TC.set_timecourse()
    TC.set_report()
    t0 = time.time()
    TC.simulate()
    ctime = time.time() - t0

    return [chem, mooseTime, htTime, htTime2, ctime, runtime ]

def main():
    ret = []
    for k, h, c, t in modelList:
        ret.append( runSim( k, h, c, t ) )

    #print("Model            MOOSE       HT_time    HT_steady_state  COPASI  simtime" )
    print( "{:18s}{:>12s}{:>12s}{:>12s}{:>12s}{:>12s}".format( "Model","MOOSE","HT_time", "HT_ss", "COPASI",  "simtime" ) )
    for model, mooseTime, htTime, htTime2, ctime, runtime  in ret:
        print( "{:18s}{:12.3f}{:12.3g}{:12.3g}{:12.3f}{:12.2g}".format( model, mooseTime, htTime/htReps, htTime2/htReps, ctime, runtime ))

if __name__ == '__main__':
    main()
```
