## Supplementary Code for "HillTau: A fast, compact abstraction for model reduction in biochemical signaling networks": fig6_plotting.py

```
# 
# This program is free software; you can redistribute it and/or
# modify it under the terms of the GNU General Public License as
# published by the Free Software Foundation; either version 3, or
# (at your option) any later version.
# 
# This program is distributed in the hope that it will be useful,
# but WITHOUT ANY WARRANTY; without even the implied warranty of
# MERCHANTABILITY or FITNESS FOR A PARTICULAR PURPOSE.  See the GNU
# General Public License for more details.
# 
# You should have received a copy of the GNU General Public License
# along with this program; see the file COPYING.  If not, write to
# the Free Software Foundation, Inc., 51 Franklin Street, Fifth
# Floor, Boston, MA 02110-1301, USA.
# 

'''
*******************************************************************
 * File:            fig6.py
 * Description:
 * Author:          Upinder S. Bhalla
 *
 ********************************************************************/

/**********************************************************************
** This program simply plots tables filled in with numbers of parameters
** and runtimes for HT and MOOSE.
**           copyright (C) 2020 Upinder S. Bhalla. and NCBS
**********************************************************************/
'''
import matplotlib.pyplot as plt
import numpy as np
from sklearn.linear_model import LinearRegression

def plotBoilerplate( panelTitle, plotPos, xlabel = 'Time (s)', ylabel = 'Conc ($\mu$M)' ):
    panelX = -0.05
    ax = plt.subplot( 3, 1, plotPos )
    ax.spines['top'].set_visible( False )
    ax.spines['right'].set_visible( False )
    '''
    ax.tick_params( direction = 'out' )
    '''
    ax.set_xlabel( xlabel, fontsize = 14 )
    ax.set_ylabel( ylabel, fontsize = 14 )
    ax.text( panelX, 1.1, panelTitle, fontsize = 18, weight = 'bold', transform = ax.transAxes )
    return ax

def regress( ax, ay, xmin, name ):
    x = np.array( ax ).reshape( ( -1, 1 ) )
    y = np.array( ay )
    model = LinearRegression()
    model.fit( x, y )
    r_sq = model.score( x, y )
    slope = model.coef_
    intcpt = model.intercept_
    xmax = max( x ) * 1.1
    print( "{}: r_sq = {}, slope = {}, intcpt = {}".format( name, r_sq, slope, intcpt ) )
    #return( [xmin, xmax], [ intcpt + slope * xmin, intcpt + slope * xmax ] )
    return( ax, [ intcpt + slope * i for i in ax ] )

def makePlots():
    massActionParams = [3,3,16,20,37,361,748]
    HillTauParams =    [3,3,5,  13,12, 31, 35]
    HillTauParams2 =    [3,3,5,  12,13, 31, 35]

    MooseRuntime = [ 1.59,1.52,3.9, 60.0, 4.9,  1520, 32000 ]

    CopasiRuntime = [ 17.9,17.3,21.0, 24.4, 26.5,   295, 4480 ]
    HillTauRuntime = [ 0.053,0.053,0.081,0.195, 0.208, 0.277, 0.68 ]
    HillTauRuntime2 = [ 0.053,0.053,0.081, 0.207, 0.195, 0.277,0.68 ]
    ax = plotBoilerplate( "A", 1, xlabel = "# Mass-action parameters", ylabel = "# HillTau params" )
    ax.set_xscale( "log" )
    ax.set_xlim( (1, 1000 ) )
    ax.set_ylim( (0, 40 ) )
    ax.plot( massActionParams , HillTauParams, "o", label = "" )
    #x, y = regress( massActionParams, HillTauParams, 1, "A" )
    #ax.plot( x, y, "b-" )

    ax = plotBoilerplate( "B", 2, xlabel = "# Mass-action parameters", ylabel = "Runtime ($\mu$s/s)" )
    ax.set_xscale( "log" )
    ax.set_yscale( "log" )
    ax.set_xlim( (1, 1000 ) )
    ax.plot( massActionParams , MooseRuntime, "-o", label = "MOOSE" )
    ax.plot( massActionParams , CopasiRuntime, ":o", label = "COPASI" )
    ax.plot( massActionParams , HillTauRuntime, "--o", label = "HillTau" )
    ax.legend( loc = 'upper left', frameon = False )
    '''
    x, y = regress( massActionParams, massActionRuntime, 3, "B MA" )
    ax.plot( x, y, "b-" )
    x, y = regress( massActionParams, HillTauRuntime, 3, "B HT" )
    ax.plot( x, y, "r-" )
    '''

    ax = plotBoilerplate( "C", 3, xlabel = "# HillTau parameters", ylabel = "Runtime ($\mu$s/s)" )
    ax.plot( HillTauParams2 , HillTauRuntime2, "o", label = "ma" )
    x, y = regress( HillTauParams2, HillTauRuntime2, 0, "C" )
    ax.set_xlim( (0, 40 ) )
    ax.set_ylim( (0, 0.7 ) )
    ax.plot( x, y, "b-" )


def main():
    fig = plt.figure( figsize = (5,7), facecolor='white' )
    fig.subplots_adjust( bottom = 0.3 )
    fig.subplots_adjust( left = 0.18 )

    makePlots()


    plt.tight_layout()
    plt.show()


if __name__ == '__main__':
    main()
```
