## Supplementary Code for "HillTau: A fast, compact abstraction for model reduction in biochemical signaling networks": hillTau.py

```
# 
# This program is free software; you can redistribute it and/or
# modify it under the terms of the GNU General Public License as
# published by the Free Software Foundation; either version 3, or
# (at your option) any later version.
# 
# This program is distributed in the hope that it will be useful,
# but WITHOUT ANY WARRANTY; without even the implied warranty of
# MERCHANTABILITY or FITNESS FOR A PARTICULAR PURPOSE.  See the GNU
# General Public License for more details.
# 
# You should have received a copy of the GNU General Public License
# along with this program; see the file COPYING.  If not, write to
# the Free Software Foundation, Inc., 51 Franklin Street, Fifth
# Floor, Boston, MA 02110-1301, USA.
# 

'''
*******************************************************************
 * File:            hillTau.py
 * Description:
 * Author:          Upinder S. Bhalla
 *
 ********************************************************************/
 '''
from __future__ import print_function
import sys
import json
import re
import argparse
import numpy as np
import matplotlib.pyplot as plt

lookupQuantityScale = { "M": 1000.0, "mM": 1.0, "uM": 1e-3, "nM": 1e-6, "pM": 1e-9 }

INTERNAL_DT_SCALE = 0.02

SIGSTR = "{:.4g}" # Used to format floats to keep to 4 sig fig. Helps when dumping JSON files.

mathFns = ["exp", "log", "ln", "log10", "abs", "sin", "cos", "tan", "sinh", "cosh", "tanh", "sqrt", "pow"]

def loadHillTau( fname ):
    with open( fname ) as json_file:
        model = json.load( json_file )
        ft = model.get( "FileType" )
        if not ft:
            print( 'Warning: Is "{}" a HillTau file? It lacks "FileType": "HillTau" specification.'.format(fname) )
        else:
            if ft != "HillTau":
                raise( ValueError( "Error: FileType should be 'HillTau', was {}.".format( ft ) ) )
    return model

def subsetModel( model, subsetInfo ):
    return

class Stim():
    def __init__( self, stim, model, off = False ):
        self.objname = stim[0]
        self.mol = model.molInfo.get( stim[0] )
        if not self.mol:
            print( "Stimulus Molecule '{}' not found".format( stim[0] ) )
            quit()
        self.value = stim[1]
        self.isOff = off
        if off:
            self.time = stim[3]
            self.value = self.mol.concInit
        else:
            self.time = stim[2]

    @staticmethod
    def stimOrder( stim ):
        return stim.time


class MolInfo():
    def __init__( self, name, grp, concInit = -1.0 ) :
        self.name = name
        self.grp = grp
        if concInit < 0.0:
            self.concInit = 0.0
            self.explicitConcInit = False
        else:
            self.concInit = concInit
            self.explicitConcInit = True
        self.order = 0  # Start out all zero
                        # First pass: 0 Reac or Eqn set to -1
                        # Second pass: cumulative order depending on subs
        self.index = 0  # Points to the conc and concInit vectors
        #print( "MolInfo {}: concInit = {}".format( name, concInit ) )

class ReacInfo():
    def __init__( self, name, grp, reacObj, molInfo, consts ):
        self.name = name
        self.grp = grp
        self._KA = convConst( consts, reacObj["KA"] )
        self.tau = convConst( consts, reacObj["tau"] )
        self.tau2 = self.tau
        self.prdIndex = molInfo[ name ].index
        molInfo[ name ].grp = grp # Put product molecule in same grp as reac
        self.subs = reacObj["subs"]
        assert( len( self.subs ) > 0 )
        if len( self.subs ) == 0:
            raise( ValueError( "Error: Reaction {} has zero reagents.".format( name ) ) )
        self.hillIndex = molInfo[ self.subs[-1] ].index
        self.reagIndex = molInfo[ self.subs[0] ].index
        self.Kmod = 1.0 # This is the default halfmax of the modifier
        self.Amod = 4.0 # This is the default alpha factor for modifier. 
        # Values of Amod < 1 make it an inhibitory modifier. 
        # Values of Amod > 1 are excitatory. Amod == 1 does not modify.
        self.Nmod = 1.0 # This is the order of the modifier.
        self.modIndex = -1

        self.gain = 1.0
        self.overrideConcInit = not molInfo[ name ].explicitConcInit
        Kmod = reacObj.get( "Kmod" )
        Amod = reacObj.get( "Amod" )
        if Amod:
            self.Amod = convConst( consts, Amod )
        Nmod = reacObj.get( "Nmod" )
        if Nmod:
            self.Nmod = convConst( consts, Nmod )
        numUnique = len( set( self.subs ) )
        self.oneSub = (numUnique == 1)
        if numUnique > 3:
            raise( ValueError( "Error: Reaction {} has > 3 reagents.".format( name ) ) )
        if numUnique == 3 and not Kmod:
            raise( ValueError( "Error: Reaction {} has 3 reagents but no Kmod specified for modifier.".format( name ) ) ) 
        if numUnique <= 2 and Kmod:
            raise( ValueError( "Error: Reaction {} has <=2 reagents, expecting 3 since Kmod specified for modifier.".format( name ) ) )
        self.HillCoeff = len( self.subs ) + 1 - numUnique
        # Hill Coeff applies only to the last mol in the list.
        self.kh = self._KA ** self.HillCoeff # Precompute it.
        self.inhibit = 0
        inh = reacObj.get( "inhibit" )
        if inh and inh != 0:
            self.inhibit = 1
        self.baseline = 0.0
        b = reacObj.get( "baseline" )
        if b:
            self.baseline = convConst( consts, b )
        tau2 = reacObj.get( "tau2" )
        if tau2:
            self.tau2 = convConst( consts, tau2 )
        if Kmod: # Various validity checks have been done above
            self.Kmod = convConst( consts, Kmod )
            self.modIndex = molInfo[ self.subs[1] ].index
        gain = reacObj.get( "gain" )
        if gain:
            self.gain = convConst( consts, gain )

        #print( "Reac {}: hillIndex={}, hillCoeff = {}, kh = {}, reagIndex  {}".format( self.name, self.hillIndex, self.HillCoeff, self.kh, self.reagIndex ) )

    def concInf( self, conc ):
        h = conc[self.hillIndex] ** self.HillCoeff
        if self.modIndex != -1:
            x = pow( conc[ self.modIndex ] / self.Kmod, self.Nmod )
            mod = ( 1.0 + x ) / ( 1.0 + self.Amod * x )
        else:
            mod = 1.0
        if self.oneSub: # this really only works for A<=>B and 2A<=>B
            return h / self._KA
        s = conc[ self.reagIndex ] * self.gain
        if self.inhibit:
            return s * (1.0 - h / ( h + self.kh * mod ) )
        else:
            return s * h / ( h + self.kh * mod )

    def concFracUp( self, t ):
        return 1.0 - np.exp( -t/self.tau )

    def concFracDown( self, t ):
        return 1.0 - np.exp( -t/self.tau2 )

    def eval( self, model, dt ):
        orig = model.conc[self.prdIndex] - self.baseline
        delta = self.concInf( model.conc ) - orig
        if delta >= 0:
            delta *= self.concFracUp( dt )
        else:
            delta *= self.concFracDown( dt )
        ret = self.baseline + orig + delta
        if ret < 0.0:
            print( "Error: negative value on: ", self.name, ret, model.conc[self.prdIndex], self.baseline, delta )
            quit()
        model.conc[self.prdIndex] = ret
        return ret
    
    def getReacField( self, field ):
        return 0.0

    @property
    def KA( self ):
        return self._KA

    @KA.setter
    def KA( self, val ):
        self._KA = val
        self.kh = self._KA ** self.HillCoeff # Precompute it.

class EqnInfo():
    def __init__( self, name, grp, eqnStr, subs, cs ):
        self.name = name
        self.grp = grp
        self.eqnStr = eqnStr
        self.subs = subs
        self.consts = cs

    def parseEqn( self, molInfo, globalConsts ):
        # Replace mol names with lookup index in molecule array, and func names with np.funcName.
        self.index = molInfo[self.name].index
        molInfo[ self.name ].grp = self.grp # Put output molecule in same grp as eqn
        self.newEq = self.eqnStr
        for name in self.consts:
            val = globalConsts.get( name )
            if val:
                self.newEq = self.newEq.replace( name, str( val ) )
            else:
                raise( ValueError( "Error: unknown const '{}' in equation '{}'".format( name, self.eqnStr ) ) )

        for mol in self.subs:
            mi = molInfo.get( mol )
            if mi:
                self.newEq = self.newEq.replace( mol, "m[{}]".format( mi.index ), 1 )
            else:
                raise( ValueError( "Error: unknown molecule '{}' in equation '{}'".format( mol, self.eqnStr ) ) )

        for func in mathFns:
            self.newEq = self.newEq.replace( func, "np.{}".format( func ) )


    def eval( self, m ):
        m[self.index] = ret = eval( self.newEq )
        return ret

class Model():
    def __init__( self, jsonDict ):
        self.jsonDict = jsonDict
        self.consts = {}
        self.molInfo = {}
        self.reacInfo = {}
        self.eqnInfo = {}
        self.grpInfo = []
        self.namedConsts = {}
        self.sortedReacInfo = []
        self.sortedEqnInfo = []
        self.currentTime = 0.0
        self.step = 0
        self.conc = np.zeros(1)
        self.concInit = np.zeros(1)
        self.plotvec = []
        self.runtime = 0.0
        self.dt = 1.0
        self.internalDt = 1.0
        self.minTau = 1.0

    '''
    def setConc( self, molName, val ):
        assert( molName in self.molInfo )
        idx = self.molInfo[ molName ].index
        self.conc[idx] = val
    '''
    def advance( self, runtime, settle = False ):
        if runtime < 10.0e-6:
            return
        if settle: 
            # This is used when we are doing a steady-state calc. Since
            # HillTau does this anyway, we jump fast. Only issue arises
            # if there are feedback processes. So to be conservative, 
            # do 10 steps. 
            innerAdvance( runtime, runtime / 10.0 )
        else:
            newdt = min( self.dt, self.internalDt )
            adv = max( self.minTau * 10.0, self.dt )
            if newdt >= runtime / 2.0:
                newdt = 10.0 ** ( np.floor( np.log10( runtime / 2.0 ) ) )
                self.innerAdvance( runtime, newdt )
            elif 2.0 * adv < runtime: 
                # Advance a few small timesteps, then switch to regular dt
                self.innerAdvance( adv, newdt )
                self.innerAdvance( runtime - adv, self.dt )
            else:   # all small dt
                self.innerAdvance( runtime, newdt )

    def innerAdvance( self, runtime, newdt ):
        # The above guarantees that newdt <= self.dt, except dose response
        t = 0.0
        while t < runtime:
            # Sometimes we have runtimes that are not a multiple of self.dt
            if newdt > ( runtime - t):
                newdt = runtime - t

            # Here we advance the simulation
            for ar in self.sortedReacInfo:
                for r in ar:
                    r.eval( self, newdt )
            for val in self.sortedEqnInfo:
                val.eval( self.conc )

            # Here we decide if we insert data into the plots.
            if np.floor( (self.currentTime + t + newdt)/ self.dt ) > self.step:
                self.step += 1
                self.plotvec.append( np.array( self.conc ) )
            t += newdt
        self.currentTime += runtime
                
    def neatRound( x ):
        if x <= 0.0:
            return 0.0
        y = pow( 10.0, np.floor( np.log10( x ) ) )
        if x/y >= 5.0:
            return y * 5.0
        elif x/y >= 2.0:
            return y * 2.0
        return y

    def reinit( self ):
        # ConcInit is evaluated only for reactions not explicitly defined.
        self.currentTime = 0
        self.step = 0
        self.internalDt = self.dt
        self.minTau = 1.0e20
        for ar in self.sortedReacInfo:
            for r in ar:
                self.minTau = min( self.minTau, r.tau, r.tau2 )
                if r.overrideConcInit:
                    if r.inhibit:
                        self.concInit[ r.prdIndex ] = r.concInf( self.concInit ) + r.baseline
                        if self.concInit[r.prdIndex] < 0.0:
                            self.concInit[r.prdIndex] = 0.0
                    else:
                        self.concInit[ r.prdIndex ] = r.baseline

        if self.dt > INTERNAL_DT_SCALE * self.minTau:
            self.internalDt = Model.neatRound( INTERNAL_DT_SCALE * self.minTau )

        # need to explicitly make new array because Python defaults to 
        # shallow copy.
        # So if you change values in conc, they will change in concInit
        self.conc = np.array( self.concInit )
        del self.plotvec[:]
        self.plotvec.append( np.array( self.conc ) )

    def getConcVec( self, molIndex ):
        return np.array( [ v[molIndex] for v in self.plotvec ] )

    def modifySched( self, saveList, deleteList ):
        numSeq = len( self.sortedReacInfo )
        newsri = [[]] * numSeq
        if len( saveList ) > 0:
            for seq in range( numSeq ):
                sri = self.sortedReacInfo[seq]
                for ri in sri:
                    if len( deleteList ) > 0 :
                        if ri in saveList and not (ri in deleteList):
                            newsri[seq].append( ri )
                    elif ri in saveList:
                        newsri[seq].append( ri )
            self.sortedReacInfo = newsri
            if len( deleteList ) > 0:
                self.sortedEqnInfo = [ val for key, val in self.eqnInfo.items() if key in saveList and not key in deleteList ]
            else: 
                self.sortedEqnInfo = [ val for key, val in self.eqnInfo.items() if key in saveList ]
        elif len( deleteList ) > 0: # Nothing on saveList
            for seq in range( numSeq ):
                sri = self.sortedReacInfo[seq]
                for ri in sri:
                    if not ( ri in deleteList ):
                        newsri[seq].append( ri )
            self.sortedReacInfo = newsri
            self.sortedEqnInfo = [ val for key, val in self.eqnInfo.items() if not key in deleteList ]
        # If both lists are empty, retain original sortedReacInfo and sortedEqnInfo.

def getQuantityScale( jsonDict ): 
    qu = jsonDict.get( "QuantityUnits" )
    qs = 1.0
    if not qu:
        qu = jsonDict.get( "quantityUnits" )
    if qu:
         qs = lookupQuantityScale[qu]

    return qs

def scaleConst( holder, name, qs, consts, constDone ):
    # This scales values with concentration units. It checks if the value
    # is a string, in which case it tries to scale the reference constant.
    # Otherwise it scales the entry in situ.
    val = holder.get( name )
    if isinstance( val, str ):
        constName = val
        ci = consts.get( constName )
        if not ci:
            raise( ValueError( "Error: Constant {} not found.".format( constName ) ) )
        if not constDone[ constName ]:
            consts[ constName] = float( SIGSTR.format( ci * qs ) )
            constDone[ constName ] = 1
    else:
        holder[name] = float( SIGSTR.format( val * qs ) )


def scaleDict( jsonDict, qs ):
    # This acts before parsing the model, so it should leave the model
    # definition layout intact. That means it should scale the
    # constants rather than fill in values where the the constants are
    # cited in the model. This means we have to take care not to scale
    # a given constant more than once, as it may be used many times.
    consts = jsonDict.get("Constants")
    if not consts:
        consts = {}
    constDone = { i:0 for i in consts.keys()} # dict of all const names.
    for grpname, grp in jsonDict["Groups"].items():
        sp = grp.get( "Species" )
        if sp:
            for m in sp:
                scaleConst( sp, m, qs, consts, constDone )
        if "Reacs" in grp:
            for reacname, reac in grp['Reacs'].items():
                # Check if it is a single substrate reac
                if len( reac["subs"] ) != 1:
                    scaleConst( reac, "KA", qs, consts, constDone )
                    #print( "REAC {} KA = {} ".format( reacname, reac["KA"] ))
                '''
                reac["tau"] = float( SIGSTR.format( reac["tau"] ) )
                tau2 = reac.get( "tau2" )
                if tau2:
                    reac["tau2"] = float( SIGSTR.format( tau2 ) )
                '''
                bl = reac.get( "baseline" )
                if bl:
                    scaleConst( reac, "baseline", qs, consts, constDone )
                kmod = reac.get( "Kmod" )
                if kmod:
                    scaleConst( reac, "Kmod", qs, consts, constDone )

def extractSubs( expr, consts ):
    # This function extracts the molecule names from a math expression.
    isInMol = 0
    molname = ""
    lastch = ''
    mols = []
    for ch in expr:
        if isInMol:
            if ch.isalnum() or ch == '_':
                molname += ch
                continue
            else:
                isInMol = 0
                if not molname in mathFns:
                    mols.append( molname )
        else:
            if ch in "eE" and (lastch.isdigit() or lastch == '.'):
                # This is the e in sci notation
                lastch = ch
                continue
            if ch.isalpha():
                molname = ch
                isInMol = 1
            else:
                lastch = ch
    if isInMol:
        mols.append( molname )
    s = []
    c = []
    for key in mols:
        if key in consts:
            c.append( key )
        else:
            s.append( key )
    return s, c

def convConst( consts, value ):
    # Convert named const to number, or if already a number, return it.
    if isinstance( value, str ):
        ret = consts.get( value )
        if ret:
            return ret
        else:
            raise( ValueError( "Error: Const {} not found.".format( value ) ) )
    return value


def parseModel( jsonDict ):
    model = Model( jsonDict )
    # First, assign all constants. Simple matter of copying the dict.
    if "Constants" in model.jsonDict:
        model.consts = model.jsonDict['Constants']
    else:
        model.consts = {}
    model.namedConsts = model.consts

    # Second, pull together all the species names. They crop up in
    # the Species, the Reacs, and the Eqns. They should be used as
    # an index to the conc and concInit vector.
    # Note that we have an ordering to decide which mol goes in which group:
    # Species; names of reacs, First term of Eqns, substrates.
    # This assumes that every quantity term has already been scaled to mM.
    for grpname, grp in model.jsonDict['Groups'].items():
        model.grpInfo.append( grpname )
        # We may have repeats in the species names as they are used 
        # in multiple places.
        if "Reacs" in grp:
            for reacname, reac in grp['Reacs'].items():
                for subname in reac["subs"]:
                    model.molInfo[subname] = MolInfo( subname, grpname)

    for grpname, grp in model.jsonDict['Groups'].items():
        if "Eqns" in grp:
            for lhs, expr in grp["Eqns"].items():
                subs, cs = extractSubs( expr, model.consts )
                for subname in subs:
                    model.molInfo[subname] = MolInfo( subname, grpname)
                model.molInfo[lhs] = MolInfo( lhs, grpname)
                model.eqnInfo[lhs] = EqnInfo( lhs, grpname, expr, subs, cs )
        if "Reacs" in grp:
            for reacname, reac in grp['Reacs'].items():
                model.molInfo[reacname] = MolInfo( reacname, grpname)

    for grpname, grp in model.jsonDict['Groups'].items():
        if "Species" in grp:
            for molname, conc in grp['Species'].items():
                model.molInfo[molname] = MolInfo( molname, grpname, concInit = convConst( model.consts, conc ) )
                grp['Species'][molname] = convConst( model.consts, conc )

    # Then assign indices to these unique molnames, and build up the
    # numpy arrays for concInit and conc.
    numMols = len( model.molInfo )
    model.conc = np.zeros( numMols )
    model.concInit = np.zeros( numMols )
    i = 0
    for molname, info in model.molInfo.items():
        info.index = i
        model.conc[i] = model.concInit[i] = info.concInit
        i += 1

    # Now set up the reactions. we need the mols all defined first.
    for grpname, grp in model.jsonDict['Groups'].items():
        if "Reacs" in grp:
            for reacname, reac in grp['Reacs'].items():
                r = ReacInfo( reacname, grpname, reac, model.molInfo, model.consts )
                model.reacInfo[reacname] = r
                # Override group so that grp of product mol == grp of reac.
                model.molInfo[ reacname ].grp = grpname
                model.molInfo[ reacname ].order = -1

    # Now set up the equation, again, we need the mols defined.
    for eqname, eqn in model.eqnInfo.items():
        eqn.parseEqn( model.molInfo, model.consts )
        # Override group so that grp of eqn output == grp of eqn.
        model.molInfo[ eqname ].grp = eqn.grp

    sortReacs( model )
    model.reinit()
    return model

def breakloop( model, maxOrder, numLoopsBroken  ):
    for reacname, reac in sorted( model.reacInfo.items() ):
        if model.molInfo[reacname].order < 0:
            model.molInfo[reacname].order = maxOrder
            #print("Warning; Reaction order loop. Breaking {} loop for {}, assigning order: {}".format( numLoopsBroken, reacname, maxOrder ) )
            return

def sortReacs( model ):
    # Go through and assign levels to the mols and reacs within a group.
    # This will be used later for deciding evaluation order.
    maxOrder = 0
    numLoopsBroken = 0
    numOrdered = 0
    numReac = len( model.reacInfo )
    while numOrdered < numReac: 
        numOrdered = 0
        stuck = True
        for reacname, reac in sorted( model.reacInfo.items() ):
            prevOrder = model.molInfo[reacname].order
            maxOrder = max( maxOrder, prevOrder )
            if prevOrder >= 0:
                numOrdered += 1
            else:
                order = [ model.molInfo[i].order for i in reac.subs ]
                if min( order ) >= 0:
                    mo = max( order ) + 1
                    model.molInfo[reacname].order = mo
                    maxOrder = max( maxOrder, mo )
                    numOrdered += 1
                    stuck = False
        #print ( "numOrdered = ", numOrdered, " / ", numReac, " max = ", maxOrder )
        if stuck:
            breakloop( model, maxOrder+1, numLoopsBroken )
            numLoopsBroken += 1

        '''
        for eqname, eqn in sorted( model.eqnInfo.items() ):
            order = [ model.molInfo[i].order for i in eqn.subs ]
            model.molInfo[eqname].order = max(order)
        '''

    maxOrder += 1
    model.sortedReacInfo = [[] for i in range( maxOrder )]
    for name, reac in model.reacInfo.items():
        order = model.molInfo[name].order
        model.sortedReacInfo[order].append( reac )
    model.sortedEqnInfo = [ val for val in model.eqnInfo.values() ]

def writeOutput( fname, model, plotvec, x ):
    with open( fname, "w" ) as fd:
        olist = sorted([ i for i in model.molInfo])
        header = "Time\t"
        outvec = [[str(v) for v in x]]
        rx = range( len( x ) )
        for name in olist:
            header += name + "\t"
            idx = model.molInfo[name].index
            outvec.append( [str(v) for v in plotvec[idx] ] )
        ry = range( len( outvec ) )
        fd.write( header + "\n" )
        for i in rx:
            for j in ry:
                fd.write( outvec[j][i] + "\t" )
            fd.write( "\n" )


def main():
    parser = argparse.ArgumentParser( description = 'This is the hillTau simulator.\n'
    'This program simulates abstract kinetic/neural models defined in the\n'
    'HillTau formalism. HillTau is an event-driven JSON form to represent\n'
    'dynamics of mass-action chemistry and neuronal activity in a fast, \n'
    'reduced form. The hillTau program loads and checks HillTau models,\n'
    'and optionally does simple stimulus specification and plotting\n')
    parser.add_argument( 'model', type = str, help='Required: filename of model, in JSON format.')
    parser.add_argument( '-r', '--runtime', type = float, help='Optional: Run time for model, in seconds. If flag is not set the model is not run and there is no display', default = 0.0 )
    parser.add_argument( '-dt', '--dt', type = float, help='Optional: Time step for model calculations, in seconds. If this argument is not set the code calculates dt to be a round number about 1/100 of runtime.', default = -1.0 )
    parser.add_argument( '-s', '--stimulus', type = str, nargs = '+', action='append', help='Optional: Deliver stimulus as follows: --stimulus molecule conc [start [stop]]. Any number of stimuli may be given, each indicated by --stimulus. By default: start = 0, stop = runtime', default = [] )
    parser.add_argument( '-p', '--plots', type = str, help='Optional: plot just the specified molecule(s). The names are specified by a comma-separated list.', default = "" )
    parser.add_argument( '-o', '--output', type = str, metavar = "fname", help='Optional: Generate an output tab-separated text file with columns of time conc1 conc2 and so on.' )
    args = parser.parse_args()
    jsonDict = loadHillTau( args.model )
    qs = getQuantityScale( jsonDict )
    scaleDict( jsonDict, qs )
    model = parseModel( jsonDict )

    runtime = args.runtime
    if runtime <= 0.0:
        return

    if args.dt < 0:
        model.dt = 10 ** (np.floor( np.log10( runtime )) - 2.0)
        if runtime / model.dt > 500:
            model.dt *= 2
    else:
        model.dt = args.dt

    stimvec = []
    
    stimMolNames = []
    for i in args.stimulus:
        if len( i ) < 2:
            print( "Warning: need at least 2 args for stimulus, got {i}".format( i ) )
            continue
        stimMolNames.append( i[0] )
        i[1] = float( i[1] ) * qs # Assume stim units same as model units.
        if len(i) == 2:
            i.extend( [0.0, runtime] )
        if len(i) == 3:
            i.extend( [runtime] )
        i[2] = float( i[2] )
        i[3] = float( i[3] )
        runtime = max( runtime, i[3] )
        stimvec.append( Stim( i, model ) )
        stimvec.append( Stim( i, model, off = True ) )

    stimvec.sort( key = Stim.stimOrder )
    model.modifySched( saveList = [], deleteList = list( set( stimMolNames )) )

    model.reinit()
    currTime = 0.0
    for s in stimvec:
        model.advance( s.time - currTime )
        model.conc[s.mol.index] = s.value
        currTime = s.time
    if runtime > currTime:
        model.advance( runtime - currTime )

    plotvec = np.transpose( np.array( model.plotvec ) )
    x = np.array( range( plotvec.shape[1] ) ) * model.dt
    clPlots = args.plots.split(',')
    if len( args.plots ) > 0 :
        clPlots = [ i.strip() for i in clPlots if i in model.molInfo]
    else: 
        clPlots = [ i for i in model.molInfo ]

    if args.output:
        writeOutput( args.output, model, plotvec, x )

    qu = jsonDict.get( "QuantityUnits" )
    if not qu:
        qu = jsonDict.get( "quantityUnits" )
    if qu:
        ylabel = 'Conc ({})'.format( qu )
        qs = lookupQuantityScale[qu]
    else:
        ylabel = 'Conc (mM)'
        qs = 1

    for name in clPlots:
        mi = model.molInfo[name]
        i = mi.index
        plt.plot( x, plotvec[i]/qs, label = name )

    plt.xlabel('Time (s)')
    plt.ylabel(ylabel)
    plt.title( args.model )
    plt.legend()
    plt.show()


if __name__ == '__main__':
    main()
```
