## Supplementary Code for "HillTau: A fast, compact abstraction for model reduction in biochemical signaling networks": ht2sbml.py

```
# 
# This program is free software; you can redistribute it and/or
# modify it under the terms of the GNU General Public License as
# published by the Free Software Foundation; either version 3, or
# (at your option) any later version.
# 
# This program is distributed in the hope that it will be useful,
# but WITHOUT ANY WARRANTY; without even the implied warranty of
# MERCHANTABILITY or FITNESS FOR A PARTICULAR PURPOSE.  See the GNU
# General Public License for more details.
# 
# You should have received a copy of the GNU General Public License
# along with this program; see the file COPYING.  If not, write to
# the Free Software Foundation, Inc., 51 Franklin Street, Fifth
# Floor, Boston, MA 02110-1301, USA.
# 

'''
*******************************************************************
 * File:            ht2sbml.py
 * Description:     Program to convert HILLTAU (JSON) files to SBML.
 * Author:          Upinder S. Bhalla
 *
 ********************************************************************/

/**********************************************************************
** This program uses HILLTAU and MOOSE and optimizes parameters of the
** HILLTAU model to fit the MOOSE one.
**           copyright (C) 2021 Upinder S. Bhalla. and NCBS
**********************************************************************/
'''
from __future__ import print_function
import sys
import os
import json
import argparse
import numpy as np
import hillTau
import simplesbml


def conv2sbml( htfile, sbmlfile, stimMol = "", events = [] ):
    # Events are a list of [time, conc] assignments.
    jsonDict = hillTau.loadHillTau( htfile )
    hillTau.scaleDict( jsonDict, hillTau.getQuantityScale( jsonDict ) )
    htmodel = hillTau.parseModel( jsonDict )

    # All 'reactants' are handled as modifiers in HillTau: they affect
    # product formation but are themselves not changed. Unfortunately
    # simplesbml does not currently support them. So I'm doing a
    # simple hack: replace all references to listOfReactants to
    # listOfModifiers in the generated SBML.

    #smodel = simplesbml.SbmlModel( sub_units = 'millimole', extent_units = 'mole' )
    # SimpleSBML doesn't let me set units to anything but mole..
    # So I'll just rescale the concs.
    smodel = simplesbml.SbmlModel()
    smodel.addCompartment( 1e-15, comp_id = 'comp' )
    for name, mol in htmodel.molInfo.items():
        smodel.addSpecies( "["+name+"]", mol.concInit *1e-3, comp = 'comp' )
    for name, reac in htmodel.reacInfo.items():
        s = reac.subs
        expr = "1.0"
        local_params = {'KA': reac.KA * 1e-3, 'tau': (reac.tau+reac.tau2)/2.0, 'n': reac.HillCoeff, 'gain': reac.gain, 'Vcomp': 1e-15}
        if len( s ) == 1:   # a --> p
            reactants = s
            expr = "Vcomp * (({0}*gain/KA)-{1})/tau".format(s[0], name )
        elif s[0] == s[-1]: # n*a --> p
            reactants = [str(len( s )) + " " + s[0]]
            expr = "Vcomp * (({0}^n * gain/KA)-{1})/tau".format(s[0], name )
        elif len( s ) == 2: # a + b --> p
            reactants = [s[0], s[1]]
            if reac.inhibit:
                expr = "Vcomp * ((gain * {0} * (1-{1}/(KA + {1})))-{2})/tau".format(s[0], s[1], name )
            else:
                expr = "Vcomp * ((gain * {0} * {1}/(KA + {1}))-{2})/tau".format(s[0], s[1], name )
        elif s[1] == s[-1]: # a + nb --> p
            reactants = [s[0], str(len( s )-1) + " " + s[1]]
            if reac.inhibit:
                expr = "Vcomp * ((gain * {0} * (1-{1}^n/(KA^n + {1}^n)))-{2})/tau".format(s[0], s[1], name )
            else:
                expr = "Vcomp * ((gain * {0} * {1}^n/(KA^n + {1}^n))-{2})/tau".format(s[0], s[1], name )
        else:               # a + mod + nb --> p
            reactants = [s[0], s[1], str(len( s )-2) + " " + s[2]]
            local_params['Kmod'] = reac.Kmod
            local_params['Amod'] = reac.Amod
            local_params['Nmod'] = reac.Nmod
            if reac.inhibit:
                expr = "Vcomp * ((gain * {0} * (1-{2}^n/((KA^n*(1+({1}/Kmod)^Nmod)/(1+Amod*(({1}/Kmod)^Nmod))) + {2}^n)))-{3})/tau".format(s[0], s[1], s[-1], name )
            else:
                expr = "Vcomp * ((gain * {0} * {2}^n/((KA^n*(1+({1}/Kmod)^Nmod)/(1+Amod*(({1}/Kmod)^Nmod))) + {2}^n))-{3})/tau".format(s[0], s[1], s[-1], name )

        smodel.addReaction( reactants, [name], expr, local_params = local_params, rxn_id="r__"+name )
    for e in events:
        smodel.addEvent( trigger='time>' + str(e[0]), assignments = {stimMol:str(e[1]) }) 

    with open( sbmlfile, 'w' ) as fd:
        # Here we get into a series of hacks to convert the rectants
        # into modifiers.
        # Hack to put in modifiers instead of reactants:
        sbmlstr = smodel.toSBML().replace( "listOfReactants", "listOfModifiers" )
        # Remove stoichometry and constant terms from modifier species refs 
        sbmlstr = sbmlstr.replace( "speciesReference", "modifierSpeciesReference" )

        sbmlstr = sbmlstr.replace( " stoichiometry=\"1\" constant=\"true\"", "" )
        sbmlstr = sbmlstr.replace( " stoichiometry=\"2\" constant=\"true\"", "" )
        sbmlstr = sbmlstr.replace( " stoichiometry=\"3\" constant=\"true\"", "" )
        sbmlstr = sbmlstr.replace( " stoichiometry=\"4\" constant=\"true\"", "" )
        sbmllist = sbmlstr.split( "listOfProducts>" )

        newstr = ""
        for s in sbmllist:
            if s[-1] == '/':    # closing line for listOfProducts
                s=s.replace( '/>', ' stoichiometry="1" constant="true"/>' )
                s=s.replace( 'modifierSpeciesReference', 'speciesReference' )
            newstr += s + "listOfProducts>"

        fd.write( newstr[:-len("listOfProducts>" )] )

    '''
    for i in stimVec:
        mi = model.molInfo.get( i.hillTauMol )
        if mi:
            inputMolIndex = mi.index
            i.molIndex = inputMolIndex
            if i.conc < 0:  # Hack to specify use of initial conc
                i.conc = mi.concInit
        else:
            raise ValueError( "Nonexistent stimulus molecule: ", i.hillTauMol )
    return Mash( model, referenceOutputs, pv, args.monitor, stimVec, jsonDict )
    '''

def main():
    parser = argparse.ArgumentParser( description = "Converts HillTau model to an approximate mass-action form in SBML." )
    parser.add_argument( "HillTauModel", type=str, help = "Required: Filepath for HillTau model in JSON format" )
    parser.add_argument( "-o", "--output", type=str, default="", metavar = "filename", help = "optional: Filepath for SBML model. Defaults to same name as HillTauModel with xml suffix" )
    args = parser.parse_args()
    if args.output == "":
        sbmlfile = args.HillTauModel.split( '/' )[-1]
        sbmlfile = sbmlfile.split( '.' )[0]
        conv2sbml( args.HillTauModel, sbmlfile + ".xml" )
    else:
        conv2sbml( args.HillTauModel, args.output )


if __name__ == '__main__':
    main()
```
