## Supplementary Code for "HillTau: A fast, compact abstraction for model reduction in biochemical signaling networks": htgraph.py

```
# 
# This program is free software; you can redistribute it and/or
# modify it under the terms of the GNU General Public License as
# published by the Free Software Foundation; either version 3, or
# (at your option) any later version.
# 
# This program is distributed in the hope that it will be useful,
# but WITHOUT ANY WARRANTY; without even the implied warranty of
# MERCHANTABILITY or FITNESS FOR A PARTICULAR PURPOSE.  See the GNU
# General Public License for more details.
# 
# You should have received a copy of the GNU General Public License
# along with this program; see the file COPYING.  If not, write to
# the Free Software Foundation, Inc., 51 Franklin Street, Fifth
# Floor, Boston, MA 02110-1301, USA.
# 

'''
*******************************************************************
 * File:            htgraph.py
 * Description:
 * Author:          G.V. Harsha Rani, Upinder S. Bhalla
 *
 ********************************************************************/

/**********************************************************************
** This program converts HILLTAU models defined in JSON format to 
** reaction diagrams. It draws on the 'dot' program for graphical layout
** of networks.
**           copyright (C) 2021 Harsha Rani, Upinder S. Bhalla. and NCBS
**********************************************************************/
'''
'''
2021
Mar 31 
sub->prd is connected with double arrow,
ligand with single arrow, 
inhibit with tee and 
modifier with diamond
legends and constant are added

Apr 7: group is added

Apr 8: eqns added with pluse and sigma 

May 4: HillTau API is called for reading json file

May 10:
added matplotlib for getting colors
validation of input file type is done
output image can be saved as png or svg

May 15: set function is remove to get Unique items
May 24: Margin for the cluster is increased from 8 to 22. 

May 31: line flags for eliminating the legend and for changing colors to bw.

June 1: added features for adjusting fontsize and height on command line

June 3: Group colors and node colors added
'''

import sys,os
sys.path.insert(1, '../../PythonCode/')
from subprocess import call
import pydot
import re
import matplotlib
from collections import OrderedDict

from hillTau import *


use_bw = False

matplotcolors = []
for name,hexno in matplotlib.colors.cnames.items():
	matplotcolors.append(name)


def countX(lst, x):
	return lst.count(x)

def unique(list1):
	output = []
	for x in list1:
		if x not in output:
			output.append(x)
	return output
	#return list(set(list1))

def checkdigit(startstringdigit,sp):
	if sp.startswith(tuple('0123456789')):
		startstringdigit[sp] = "s"+sp
		
def checkdigitEqu(statstringdigit,sp):
	if sp in startstringdigit:
		sp = startstringdigit[sp]
	return(sp)

def getColor(gIndex,fwd_rev="forward"):
	if use_bw:
		return( "black", gIndex )

	if gIndex < len(matplotcolors):
		grpcolor = matplotcolors[gIndex]
		if grpcolor in ["white","wheat","aqua","whitesmoke","mintcream","oldlace","black","snow","aliceblue","azure","cornsilk","beige","bisque","blanchedalmond","antiquewhite","lightyellow","lightsteelblue","ghostwhite","floralwhite","ivory","honeydew"]:#mediumpurple","mediumvioletred","mediumseagreen"]:
			if fwd_rev == "reverse":
				gIndex = gIndex -1
			else:
				gIndex = gIndex +1

			return getColor(gIndex,fwd_rev)
		else:
			if fwd_rev == "reverse":
				gIndex = gIndex -1
			else:
				gIndex = gIndex +1
			return(grpcolor,gIndex)
	else:
		return getColor(0)

def jsontoPng(modelpath, outputfile, ranksep = 0, hasLegend = True, fontsize = 18, showGroups = True ):
	group_no = 0;
	#groupmap = dict()
	groupmap = OrderedDict()
	global startstringdigit
	startstringdigit= {}
	global node_color
	node_color = {}
	lig_exist = False
	kmod_exist = False
	inhibit_exist = False
	edge_arrowsize = 1.5
	edge_weight = 1

	s = ""
	st = os.path.splitext(outputfile)
	outputfilename = st[0]
	if len( st ) > 1: 
		outputfiletype = st[1][1:]
	else:
		outputfiletype = "png"
	f_graph = open(outputfilename+".dot", "w")
	f_graph.write("digraph mygraph {\n\trank=TB;\n")
	if ranksep > 0.0:
		f_graph.write("\tranksep={};\n".format( ranksep ))
	#f_graph.write("ratio = 1.0\n")
	#f_graph.write("ratio = \"fill\"\n")
	#f_graph.write("size = \"4,4!\"\n")
	#f_graph.write("node [shape=box, penwidth=2, height=0.01, width=0.01 ];")
	f_graph.write("node [shape=box, penwidth=2,fontsize={}];".format( fontsize ) )
	
	specielist,node_color = writeSpecies(modelpath,groupmap,f_graph)
	funclist = writeFunc(modelpath,groupmap,f_graph,edge_arrowsize,edge_weight, fontsize = fontsize - 2)
	edgelist,node_color,lig_exist,kmod_exist,inhibit_exist = writeReac(modelpath,groupmap,f_graph,edge_arrowsize,edge_weight, fontsize = fontsize - 2)
	nIndex = len(matplotcolors)-1
	if showGroups:
		for grp,items in groupmap.items():
			color,nIndex = getColor(nIndex,"reverse")
			s = s + "\nsubgraph cluster_"+str(group_no)+"i\n{"
			s = s+"\nsubgraph cluster_"+str(group_no)+"\n{"+"\n"+"label=\""+grp+"\";\npenwidth=4; margin=10.0\ncolor=\""+color+"\";\nfontsize="+str(fontsize + 2)+";\n"
			sps = ""
			items = list(unique(items))
			for sp in items:
				if items.index(sp) != 0:
					sps = sps+','+sp
				else:
					sps = sps+sp
			s = s+sps+"\n} style=invisible\n}"
			group_no += 1;
	
	f_graph.write(s)
	f_graph.write(edgelist)
	f_graph.write(funclist)
	nodeIndex = 0
	for k,v in node_color.items():
		v,nodeIndex = getColor(nodeIndex)
		f_graph.write("\n"+k+"[color=\""+v+"\"]")
	for p,q in startstringdigit.items():
		f_graph.write("\n"+q+"[label=\""+p+"\"]")	
	
	if hasLegend:
	    f_graph.write("\nnode [shape=plaintext]\nsubgraph cluster_01 {\n\tlabel = \"Legend\";\n\t{ rank=sink;\n\tkey [label=<<table border=\"0\" cellpadding=\"2\" cellspacing=\"0\" cellborder=\"0\">\n\t<tr><td align=\"right\" port=\"i1\">Input</td></tr>\n")
	    if lig_exist:
		    f_graph.write("\t<tr><td align=\"right\" port=\"i2\">Activate</td></tr>\n")
	    if kmod_exist:
		    f_graph.write("\t<tr><td align=\"right\" port=\"i3\">Modifier</td></tr>\n")
	    if inhibit_exist:
		    f_graph.write("\t<tr><td align=\"right\" port=\"i4\">Inhibit</td></tr>\n")
	    f_graph.write("\t</table>>]\n\tkey2 [label=<<table border=\"0\" cellpadding=\"2\" cellspacing=\"0\" cellborder=\"0\">\n\t<tr><td port=\"i1\">&nbsp;</td></tr>\n")
	    if lig_exist:
		    f_graph.write("\t<tr><td port=\"i2\">&nbsp;</td></tr>\n")
	    if kmod_exist:
		    f_graph.write("\t<tr><td port=\"i3\">&nbsp;</td></tr>\n")
	    if inhibit_exist:
		    f_graph.write("\t<tr><td port=\"i4\">&nbsp;</td></tr>\n")
	    f_graph.write("\t</table>>]\n\tkey:i1:e -> key2:i1:w [arrowhead=normal color=\"black:black\" style=bold]\n")
	    if lig_exist:
		    f_graph.write("\tkey:i2:e -> key2:i2:w [arrowhead=vee]\n")
	    if kmod_exist:
		    f_graph.write("\tkey:i3:e -> key2:i3:w [arrowhead=odiamond]\n")
	    if inhibit_exist:
		    f_graph.write("\tkey:i4:e -> key2:i4:w [arrowhead=tee]\n")
	    f_graph.write("\t}\n\t}")

	f_graph.write("\n}")
	f_graph.close()
	
	command = "dot -T"+ outputfiletype + " "+ outputfilename+".dot -o "+outputfile
	call([command], shell=True)

def writeSpecies(modelpath, groupmap,f_graph):
	# getting all the species
	specieslist = ""
	mIndex = 0 
	for molname, mol in ( modelpath.molInfo.items() ):
		checkdigit(startstringdigit,molname)
		molname = checkdigitEqu(startstringdigit,molname)
		if molname not in node_color:
			spe_color,mIndex = getColor(mIndex)
			node_color[molname] = spe_color

		if mol.grp in groupmap:
			groupmap[mol.grp].append(molname)
		else:
			groupmap[mol.grp] = [molname]
	return specieslist,node_color
		
def writeFunc(modelpath,groupmap,f_graph,edge_arrowsize,edge_weight, fontsize = 16):
	equation_pluse = 0
	equation_sigma = 0
	edgelist = ""
	constants_list = []
	for e,t in modelpath.eqnInfo.items():
		checkdigit(startstringdigit,t.name)
		t.name = checkdigitEqu(startstringdigit,t.name)
		allpluse = True
		mathSym = []
		for i in t.eqnStr:
			if i in ["*","-","/","+"]:
				mathSym.append(i)
		if len(unique(mathSym)) == 1 and mathSym[0] == "+":
			allpluse = True
		else:
			allpluse = False
		if allpluse:
			plusesize = "pluse"+str(equation_pluse)
			equation_pluse+=1
			edgelist = edgelist+"\n"+plusesize+"[label=\"+\",shape=circle]"
			groupmap[t.grp].append(plusesize)
		else:
			plusesize = "sigma"+str(equation_sigma)
			equation_sigma+=1
			edgelist = edgelist+"\n"+plusesize+"[label=<&Sigma;>,shape=circle]"
			groupmap[t.grp].append(plusesize)
		for tsubs in unique(t.subs):
			input_color = node_color[tsubs]
			c = countX(t.subs,tsubs)
			edgelist = edgelist+"\n"+tsubs+"->"+plusesize+"[arrowhead=vee weight = "+str(edge_weight)+" color=\""+input_color+ "\" arrowsize = "+str(edge_arrowsize)+""
			if c > 1:
				edgelist = edgelist+ " label=\" "+str(c)+"\" fontsize={}".format( fontsize )
			edgelist = edgelist+"]"
		edgelist = edgelist+"\n"+plusesize+"->"+t.name+"[arrowhead=vee weight = "+str(edge_weight)+ " arrowsize = "+str(edge_arrowsize)+"]"
	return edgelist

def writeReac(modelpath,groupmap,f_graph,edge_arrowsize,edge_weight,fontsize = 16):
	edgelist = ""
	reaction_color_list =[]
	lig_exist = False
	kmod_exist = False
	inhibit_exist = False
	sIndex = 0
	for reacname, reac in ( modelpath.reacInfo.items() ):
		checkdigit(startstringdigit,reacname)
		reacname = checkdigitEqu(startstringdigit,reacname)
		if reac.grp in groupmap:
			groupmap[reac.grp].append(reacname)
		else:
			groupmap[reac.grp] = [reacname]
		sublist = reac.subs
		sublistU = unique(reac.subs)
		prd = reacname
		#reaction_color = getRandcolorX(reaction_color_list)
		#reaction_color,sIndex = getColor(sIndex)
		#reaction_color = getRandColor()
		for sub in sublistU:
			newsub = sub
			if sub in startstringdigit:
				newsub = startstringdigit[sub]
			''' if string starting with number, then replace with s+string'''		
			if newsub in node_color:
				reaction_color = node_color[newsub]
			else:
				reaction_color,sIndex = getColor(sIndex)
				node_color[newsub] = reaction_color

			checkdigit(startstringdigit,sub)
			if reac.inhibit == 1.0 and sublist.index(sub) == 1:
				c = countX(sublist,sub)
				sub = checkdigitEqu(startstringdigit,sub)
				inhibit_exist = True
				''' inhibit  tee  '''
				if c >1:
					edgelist = edgelist+"\n"+sub+"->"+prd+"[arrowhead = tee weight = "+str(edge_weight)+ " arrowsize = "+str(edge_arrowsize)+" color=\""+reaction_color+"\" label=\" "+str(c)+"\" fontsize="+str(fontsize)+ "]"
				else:
					edgelist = edgelist+"\n"+sub+"->"+prd+"[arrowhead = tee weight = "+str(edge_weight)+ " arrowsize = "+str(edge_arrowsize)+" color=\""+reaction_color+"\"]"
			elif len(sublistU) == 3 and sublist.index(sub) == 1:
				''' kmod odiamond '''
				sub = checkdigitEqu(startstringdigit,sub)
				edgelist = edgelist+"\n"+sub+"->"+prd+"[arrowhead = odiamond weight = "+str(edge_weight)+ " arrowsize = "+str(edge_arrowsize)+" color=\""+reaction_color+"\"]"
				kmod_exist = True
			else:
				if sublist.index(sub) >= 1:
					c = countX(sublist,sub)
					lig_exist = True
					sub = checkdigitEqu(startstringdigit,sub)
					''' ligand vee '''
					if c >1:
						edgelist = edgelist+"\n"+sub+"->"+prd+"[arrowhead=vee weight = "+str(edge_weight)+ " arrowsize = "+str(edge_arrowsize)+" color=\""+reaction_color+"\" label=\" "+str(c)+"\" fontsize="+str(fontsize)+ "]"
					else:
						edgelist = edgelist+"\n"+sub+"->"+prd+"[arrowhead=vee weight = "+str(edge_weight)+ " arrowsize = "+str(edge_arrowsize)+" color=\""+reaction_color+"\"]"				
				else:
					if  sublist.index(sub) == 0:
						sub = checkdigitEqu(startstringdigit,sub)
						edgelist = edgelist +"\n"+sub+"->"+prd+"[arrowhead=normal weight = "+str(edge_weight)+ " arrowsize = "+str(edge_arrowsize)+" color=\""+reaction_color+":"+reaction_color+"\" style=bold]"				
	return(edgelist,node_color,lig_exist,kmod_exist,inhibit_exist)

def file_choices(choices,fname,iotype):
	ext = (os.path.splitext(fname)[1][1:]).lower()
	if iotype == "outputfile":
		if ext not in choices:
			parser.error("Requires output filetype {}".format(choices))
	else:
		if ext != "json":
			parser.error("Requires HillTau file in JSON format ")
			
	return fname

if __name__ == "__main__":

	parser = argparse.ArgumentParser( description = 'This program generates a reaction diagram for a HillTau model. It converts the specified HillTau file in JSON format, to the dot format. The dot file is further converted to an image in png/svg format\n')
	parser.add_argument('model',type=lambda s:file_choices(("json"),s,"input"), help='Required: filename of model, in JSON format.')
	#parser.add_argument( 'model', type = str, help='Required: filename of model, in JSON format.')
	parser.add_argument( '-o', '--output', type=lambda out:file_choices(("png","svg"),out,"outputfile"), help='Optional: writes out the png model into named file. default takes json filename')
	parser.add_argument( '-r', '--ranksep', type=float, default = 0, help='Optional: set rank separation (vertical spacing) in output.')
	parser.add_argument( '-fs', '--fontsize', type=float, default = 18, help='Optional: set font size for node labels.')
	parser.add_argument( '-nl', '--no_legend', action='store_true', help='Optional: Turns off generation of legend')
	parser.add_argument( '-ng', '--no_groups', action='store_true', help='Optional: Removes grouping. All molecules and reactions sit together.')
	parser.add_argument( '-bw', '--bw', action='store_true', help='Optional: Goes into black-and-white plotting mode')
	args = parser.parse_args()
	use_bw = args.bw

	if args.output == None:
		dirpath = os.path.dirname(args.model)
		basename = os.path.basename(args.model)
		if dirpath:
			outputfile = os.path.dirname(args.model)+"/"+os.path.splitext(os.path.basename(args.model))[0]+".png"	
		else:
			outputfile = os.path.splitext(args.model)[0]+".png"
	else:
		outputfile = args.output

	jsonDict = loadHillTau( args.model )
	modelpath = parseModel( jsonDict )
	jsontoPng(modelpath, outputfile, ranksep = args.ranksep, hasLegend = not args.no_legend, fontsize = args.fontsize, showGroups = not args.no_groups )
```
