## Supplementary Code for "HillTau: A fast, compact abstraction for model reduction in biochemical signaling networks": insert_benchmark_events.py

```
# -*- coding: utf-8 -*-
"""
Created on Tue Feb 10 15:26:54 2015
@author: carolc24
"""

import simplesbml

#with open( 'SBML_MODELS/acc92_fixed.xml', 'r' ) as fr:
with open( 'SBML_MODELS/acc92_for_COPASI.xml', 'r' ) as fr:
    modelstr = fr.read()
#modelstr = loadFromFile( 'SBML_MODELS/acc92_fixed.xml' )
model = simplesbml.loadSBMLStr( modelstr )

#model = simplesbml.loadSBMLFile( 'SBML_MODELS/acc92_fixed.xml' )

model.addEvent(trigger='time>6600', assignments={'Ca_2327_0_':'1.0e-3'}); 
model.addEvent(trigger='time>7500', assignments={'Ca_2327_0_':'0.08e-3'});
model.addEvent(trigger='time>12000', assignments={'Ca_2327_0_':'10e-3'});
model.addEvent(trigger='time>12001', assignments={'Ca_2327_0_':'0.08e-3'});
model.addEvent(trigger='time>12300', assignments={'Ca_2327_0_':'10e-3'});
model.addEvent(trigger='time>12301', assignments={'Ca_2327_0_':'0.08e-3'});
model.addEvent(trigger='time>12600', assignments={'Ca_2327_0_':'10e-3'});
model.addEvent(trigger='time>12601', assignments={'Ca_2327_0_':'0.08e-3'});

model.addEvent(trigger='time>6600', assignments={'BDNF_2052_0_':'3.7e-6'}); 
model.addEvent(trigger='time>7500', assignments={'BDNF_2052_0_':'0.05e-6'});
model.addEvent(trigger='time>12000', assignments={'BDNF_2052_0_':'3.7e-6'});
model.addEvent(trigger='time>12005',assignments={'BDNF_2052_0_':'0.05e-6'});
model.addEvent(trigger='time>12300', assignments={'BDNF_2052_0_':'3.7e-6'});
model.addEvent(trigger='time>12305',assignments={'BDNF_2052_0_':'0.05e-6'});
model.addEvent(trigger='time>12600', assignments={'BDNF_2052_0_':'3.7e-6'});
model.addEvent(trigger='time>12605',assignments={'BDNF_2052_0_':'0.05e-6'});

#code1 = simplesbml.writeCode(example.document); #Produces code to create model 'example' in string format
with open( "benchmark_acc92.xml", 'w' ) as fd:
        fd.write( model.toSBML() )
```
