## Supplementary Code for "HillTau: A fast, compact abstraction for model reduction in biochemical signaling networks": mash.py

```
# 
# This program is free software; you can redistribute it and/or
# modify it under the terms of the GNU General Public License as
# published by the Free Software Foundation; either version 3, or
# (at your option) any later version.
# 
# This program is distributed in the hope that it will be useful,
# but WITHOUT ANY WARRANTY; without even the implied warranty of
# MERCHANTABILITY or FITNESS FOR A PARTICULAR PURPOSE.  See the GNU
# General Public License for more details.
# 
# You should have received a copy of the GNU General Public License
# along with this program; see the file COPYING.  If not, write to
# the Free Software Foundation, Inc., 51 Franklin Street, Fifth
# Floor, Boston, MA 02110-1301, USA.
# 

'''
*******************************************************************
 * File:            mash2.py
 * Description:
 * Author:          Upinder S. Bhalla
 *
 ********************************************************************/

/**********************************************************************
** This program uses HILLTAU and MOOSE and optimizes parameters of the
** HILLTAU model to fit the MOOSE one.
**           copyright (C) 2021 Upinder S. Bhalla. and NCBS
**********************************************************************/
'''
from __future__ import print_function
import sys
import os
from scipy.optimize import minimize
import json
import time
import argparse
import numpy as np
import matplotlib.pyplot as plt
import moose
import moose.model_utils as mu
import hillTau

t1 = 20
t2 = 60
t3 = 100
i1 = 1e-3

plotDt = 1
stimVec = [[0, 0.0, 20.0], [0, 1e-3, 40.0], [0, 0, 40.0]]
stimRange = [ 0.1, 0.2, 0.5, 1, 2.0, 5.0, 10.0 ]
settleTimeScale = stimRange[-1]  # How much longer is settleTime than midTime?

class Stim:
    ### Advance to specified time, and then set the conc to the stim value.
    def __init__( self, mol, conc, time ):
        self.mooseMol = getMooseName( mol )
        self.hillTauMol = getHillTauName( mol )
        self.conc = conc
        self.time = time
        self.molIndex = 0

class Mash:
    def __init__( self, model, reference, params, outputMolNames, stimVec, jsonDict ):
        self.model = model
        self.reference = reference
        self.params = params
        htNames = [ getHillTauName( i ) for i in outputMolNames ]
        self.plotnum = { i:model.molInfo[ i ].index for i in htNames }
        self.stimVec = stimVec
        self.jsonDict = jsonDict
        self.numIter = 0
        self.simt = 0
        self.molMap = { getMooseName(i):getHillTauName(i) for i in outputMolNames }

    def scaleParams( self, x ):
        for i, scaleFactor in zip( self.params, x ):
            spl = i.rsplit( '.' ,1)
            assert( len(spl) == 2 )
            obj, field = spl
            if field == "concInit" or field == "conc":
                mi = self.model.molInfo[ obj ]
                mi.concInit *= scaleFactor
                self.model.concInit[ mi.index ] *= scaleFactor
            elif field == "tau": 
                # handle implicit assignment of tau2 when they are equal.
                ri = self.model.reacInfo[ obj ]
                if np.isclose( ri.tau, ri.tau2 ):
                    ri.tau = ri.tau2 = ri.tau * scaleFactor
                else:
                    ri.tau = ri.tau * scaleFactor
            else:
                orig = getattr( self.model.reacInfo[ obj ], field )
                setattr( self.model.reacInfo[ obj ], field, orig * scaleFactor )

    def doRun(self, x ):
        self.scaleParams( x )
        t0 = time.time()
        self.model.reinit()
        lastt = 0.0
        for stim in self.stimVec:
            self.model.advance( stim.time - lastt )
            self.model.conc[ stim.molIndex ] = stim.conc
            lastt = stim.time
        self.simt += time.time() - t0
        #nt = np.transpose( np.array( self.model.plotvec ) )
        #ret = { name:nt[index] for name, index in self.plotnum.items() }
        ret = { name:self.model.getConcVec( index ) for name, index in self.plotnum.items() }
        self.scaleParams( 1.0/x )
        self.numIter += 1
        return ret

    def doScore( self, outDict ):
        sq = 0.0
        for name, ref in self.reference.items():
            yrange = max( ref )
            y = ( outDict[self.molMap[name]] - ref ) / yrange
            sq += np.dot( y, y ) / len( ref )
        return np.sqrt( sq )

    def doEval( self, x ):
        ret = self.doRun( x )
        return self.doScore( ret )

    def dumpScaledFile( self, x, fname ):
        # This is significantly more complicated because the values may
        # be specified in the Constants section of the file.
        jd = self.jsonDict
        consts = jd.get( "Constants", {} )
        # Scale back to original units
        hillTau.scaleDict( jd, 1.0/hillTau.getQuantityScale( jd ) )
        # Scale each of the parameters
        for i, scaleFactor in zip( self.params, x ):
            spl = i.rsplit( '.' ,1)
            assert( len(spl) == 2 )
            obj, field = spl
            if field == "concInit":
                mi = self.model.molInfo[ obj ]
                concInit = jd["Groups"][ mi.grp ][ "Species" ][ mi.name ] 
                if isinstance( concInit, str ):
                    if not concInit in consts:
                        raise( ValueError( "Error: Constant {} not found".format( concInit ) ) )
                    else:
                        consts[ concInit ] *= scaleFactor
                else:
                    jd["Groups"][ mi.grp ][ "Species" ][ mi.name ] = concInit * scaleFactor
            else:
                ri = self.model.reacInfo[ obj ]
                orig = jd["Groups"][ ri.grp ][ "Reacs" ][obj][field]
                if isinstance( orig, str ):
                    if not orig in consts:
                        raise( ValueError( "Error: Constant {} not found".format( concInit ) ) )
                    else:
                        consts[ orig ] *= scaleFactor
                else:
                    jd["Groups"][ ri.grp ]["Reacs"][obj][field] = orig * scaleFactor

        with open( fname, 'w' ) as f:
            json.dump( jd, f, indent = 4 )

# Callback function for minimizer. Just prints out dots.
def dotter( xk ):
    print( ".", end = "", flush = True )


def makeMash( args, stimVec, referenceOutputs ):
    #jsonDict = hillTau.loadHillTau( "HT_MODELS/opt_fb_inhib.json" )
    jsonDict = hillTau.loadHillTau( args.HillTauModel )

    if len( args.addParams ) > 0:
        pv = args.addParams
    else:
        pv = paramVec( jsonDict )
    for i in args.removeParams:
        if i in pv:
            pv.remove( i )

    hillTau.scaleDict( jsonDict, hillTau.getQuantityScale( jsonDict ) )
    model = hillTau.parseModel( jsonDict )
    model.dt = plotDt
    for i in stimVec:
        mi = model.molInfo.get( i.hillTauMol )
        if mi:
            inputMolIndex = mi.index
            i.molIndex = inputMolIndex
            if i.conc < 0:  # Hack to specify use of initial conc
                i.conc = mi.concInit
        else:
            raise ValueError( "Nonexistent stimulus molecule: ", i.hillTauMol )
    return Mash( model, referenceOutputs, pv, args.monitor, stimVec, jsonDict )

def plotBoilerplate( xlabel = 'Time (s)', ylabel = 'Conc ($\mu$M)', title = "" ):
    ax = plt.subplot( 1, 1, 1 )
    ax.spines['top'].set_visible( False )
    ax.spines['right'].set_visible( False )
    ax.set_xlabel( xlabel, fontsize = 14 )
    ax.set_ylabel( ylabel, fontsize = 14 )
    ax.set_title( title )
    return ax

def runMoose( chem, stimVec, outMols ):
    filename, file_extension = os.path.splitext(chem)
    if file_extension == ".g":
        modelId = moose.loadModel( chem, 'model', 'gsl' )
    elif file_extension == ".xml":
        #modelId = mu.mooseReadSBML( chem, 'model', 'gsl' )
        modelId = moose.readSBML( chem, 'model', 'gsl' )
    '''
    moose.le( "/model/kinetics" )
    for i in moose.wildcardFind ( "/model/kinetics/##[ISA=PoolBase]" ):
        print( i.name, i.concInit )
    for i in moose.wildcardFind ( "/model/kinetics/##[ISA=Reac]" ):
        print( i.name, i.Kf, i.Kb )
    '''
    tabs = moose.Neutral( "/model/tabs" )
    mooseMols = [ getMooseName( i ) for i in outMols ]
    for i in mooseMols:
        el = moose.wildcardFind( "/model/kinetics/" + i + ",/model/kinetics/##/" + i )
        if len( el ) > 0:
            # Make an output table
            tab = moose.Table2( "/model/tabs/" + i )
            moose.connect( tab, "requestOut", el[0], "getConc" )
    for i in range( 10, 20 ):
        moose.setClock( i, plotDt )

    moose.reinit()
    lastt = 0.0

    for stim in stimVec:
        #print( "STIM = ", stim.mol, "   ", stim.conc, " ", stim.time )
        el = moose.wildcardFind( "/model/kinetics/" + stim.mooseMol + ",/model/kinetics/##/" + stim.mooseMol )
        if len( el ) > 0:
            if stim.time > lastt:
                moose.start( stim.time - lastt )
                lastt = stim.time
            el[0].concInit = stim.conc # assign conc even if no sim advance
        else:
            print( "Warning: Stimulus molecule '{}' not found in MOOSE".format( stim.mooseMol ) )

    vecs = { i.name:i.vector for i in moose.wildcardFind("/model/tabs/#") }
    return vecs

def paramVec( jsonDict ):
    pv = []
    for grp in jsonDict['Groups'].values():
        reacDict = grp.get( 'Reacs' )
        if not reacDict:
            continue
        for reacname, reac in reacDict.items():
            pv.append( reacname + ".KA" )
            pv.append( reacname + ".tau")
            tau2 = reac.get( "tau2" )
            if tau2:
                pv.append( reacname + ".tau2")
            gain = reac.get( "gain" )
            if gain:
                pv.append( reacname + ".gain")
            baseline = reac.get( "baseline" )
            if baseline:
                pv.append( reacname + ".baseline")
            Kmod = reac.get( "Kmod" )
            if Kmod:
                pv.append( reacname + ".Kmod")
            Amod = reac.get( "Amod" )
            if Amod:
                pv.append( reacname + ".Amod")

        if 'Species' in grp:
            for molname, mol in grp['Species'].items():
                pv.append( molname + ".concInit")

    return pv

def parseDoser( stimVec, d, t ):
    assert( len(d) == 3 )
    mol, midconc, settleTime = d
    midconc = float( midconc )
    settleTime = float( settleTime )
    #print("'{}'     '{}'     '{}'".format( mol, midconc, settleTime) )
    # Build dose=response
    stimVec.append( Stim( mol, 0.0, t ) )
    t += settleTime
    for x in stimRange: 
        stimVec.append( Stim( mol, midconc * x, t ) )
        t += settleTime
    stimVec.append( Stim( mol, 0.0, t ) ) 
    t += settleTime
    return t

def parseCycle( stimVec, c, t ):
    mol = c[0]
    conc, onTime, offTime = [ float( x ) for x in c[1:4] ]
    numCycles = int( c[4] )
    stimVec.append( Stim( mol, 0.0, t ) )
    for i in range( numCycles ):
        t += float( offTime )
        stimVec.append( Stim( mol, float( conc ), t ) )
        t += float( onTime )
        stimVec.append( Stim( mol, 0.0, t ) )
    t += float( offTime ) # final zero level stim
    stimVec.append( Stim( mol, 0.0, t ) )
    return t

def parseStims( stimArg, builtin, cyclic, doser ):
    stimVec = []
    t = 0.0
    for b in builtin:
        assert( len(b) == 3 ) # molecule, midconc, midTime
        mol, midconc, midTime = b
        midconc = float( midconc )
        midTime = float( midTime )
        #print("'{}'     '{}'        '{}'".format( mol, midconc, midTime) )
        settleTime = midTime * settleTimeScale
        # Build dose=response
        t = parseDoser( stimVec, [mol, midconc, settleTime], t)
        # Build cyclic stimulus
        sr0 = stimRange[0]
        t = parseCycle( stimVec, [mol, midconc, midTime*sr0, midTime*sr0, len(stimRange)*25 ], t)
        t = parseCycle( stimVec, [mol, midconc, midTime, midTime, int( len(stimRange) * 2.5 ) ], t)

    for c in cyclic:
        assert( len(c) == 5 ) # molecule, conc, start, stop, numCycles
        t = parseCycle( stimVec, c, t )

    for d in doser:
        assert( len(d) == 3 ) # molecule, midconc, settleTime
        t = parseDoser( stimVec, d, t )

    for s in stimArg:
        assert( len( s ) >= 3 and len(s) % 2 == 1 )
        for i in range( 1, len( s ), 2 ):
            stimVec.append( Stim( s[0], float( s[i] ), float(s[i+1]) ) )
    return sorted( stimVec, key = lambda x: x.time )

def oldparseStims( stimArg, builtin, cyclic, doser ):
    stimVec = []
    t = 0.0
    for b in builtin:
        # We do the dose-response, then high-freq, then mid-freq.
        # The dose response internally also does low-freq.
        assert( len(b) == 3 ) # molecule, midconc, midtime
        mol, midconc, midtime = b
        midconc = float( midconc )
        midtime = float( midtime )
        #print("'{}'     '{}'        '{}'".format( mol, midconc, midtime) )
        settleTime = midtime * settleTimeScale
        # Build dose=response
        stimVec.append( Stim( mol, 0.0, t ) )
        t += settleTime
        for x in stimRange: 
            stimVec.append( Stim( mol, midconc * x, t ) )
            t += settleTime
        # Use -ve conc to tell it to look up initial conc.
        stimVec.append( Stim( mol, -1.0, t ) ) 
        t += settleTime

        # duration, hence optimization weight, of each stim should match.
        for x in range( int( 0.25 * settleTimeScale * len( stimRange ) / stimRange[0] ) ):
            t += midtime * stimRange[0]
            stimVec.append( Stim( mol, midconc, t ) )
            t += midtime * stimRange[0]
            stimVec.append( Stim( mol, 0, t ) )

        for x in range( int( 0.25 * settleTimeScale * len( stimRange ) ) ):
            t += midtime
            stimVec.append( Stim( mol, midconc, t ) )
            t += midtime
            stimVec.append( Stim( mol, 0, t ) )
        t += midtime
        # Use -ve conc to tell it to look up initial conc.
        stimVec.append( Stim( mol, -1.0, t ) )

    for c in cyclic:
        for i in range( c[4] ):
            t += float( c[2] )
            stimVec.append( Stim( c[0], float( c[1] ), t ) )
            t += float( c[3] )
            stimVec.append( Stim( c[0], 0, t ) )
        t += float( c[2] ) # final zero level stim
        stimVec.append( Stim( c[0], float( c[1] ), t ) )

    for s in stimArg:
        assert( len( s ) >= 3 and len(s) % 2 == 1 )
        for i in range( 1, len( s ), 2 ):
            stimVec.append( Stim( s[0], float( s[i] ), float(s[i+1]) ) )
    return sorted( stimVec, key = lambda x: x.time )

def getMooseName( name ):
    sp = name.split( ':' )
    return sp[0]

def getHillTauName( name ):
    sp = name.split( ':' )
    if len(sp ) == 1:
        return name
    else:
        return sp[1]

def main():
    global plotDt
    stimDict = {}
    parser = argparse.ArgumentParser( description = "Optimizes HillTau models to fit chemical kinetic (mass action and Michaelis-Menten) models of chemical signalling." )
    parser.add_argument( "chemModel", type = str, help = "Required: Filepath for chemical kinetic model" )
    parser.add_argument( "HillTauModel", type=str, help = "Required: Filepath for HillTau model" )
    parser.add_argument( "-m", "--monitor", type = str, nargs = '+', metavar = "molName", help = "Optional: Molecules to monitor, as a list of space-separated names. If names differ between chemical and HillTau models, both can be specified, separated by a colon. Example: Ca:Calcium.", default = ["output"] )
    parser.add_argument( '-b', '--builtin', nargs = 3, metavar = ('molecule', 'midconc', 'midtime'), action='append', help='Optional: Deliver builtin stimulus. This is a dose-response centered around midconc, with a settling time of midtime * 10. This is followed by a timeseries of square-wave pulses from 0 to midconc, with on-time of midtime/10 followed by another with on-time of midtime. The first runs for 170 cycles and the second for 17. If multiple builtin stimuli are specified, they will be executed in order, without overlap. If molecule names are different between chem and HillTau models, they should be separated by a colon.', default = [] )
    parser.add_argument( '-c', '--cyclic', nargs = 5, metavar = ('molecule', 'conc', 'onTime', 'offTime', 'num_cycles'), action='append', help='Optional: Deliver cyclic stimulus. This is a timeseries of rectangular pulses from 0 to conc, with an onTime and offTime as specified, repeated for num_cycles. Before the first cycle, and after the last cycle it runs for another "offTime" seconds at conc = 0. If molecule names are different between chem and HillTau models, they should be separated by a colon.', default = [] )
    parser.add_argument( '-d', '--dose_response', nargs = 3, metavar = ('molecule', 'midconc', 'settle_time'), action='append', help='Optional: Deliver dose-response stimulus centered around midconc, with a settling time of settle_time. If other builtin, cyclic or dose_response stimuli are specified, they will be executed in order, without overlap. If molecule names are different between chem and HillTau models, they should be separated by a colon.', default = [] )
    parser.add_argument( "-a", "--addParams", type = str, nargs = "+", metavar = "obj.field", help = "Optional: Add parameter list. This will remove all the automatic ones obtained by scanning through the model, and only use the added ones from this list. Each parameter is of the form object.field. Any number of parameters can be added, separated by spaces. If molecule names are different between chem and HillTau models, they should be separated by a colon", default = [] ) 
    parser.add_argument( "-r", "--removeParams", type = str, nargs = "+", metavar = "param", help= "Optional: Remove parameters from the default ones which were generated automatically by scanning all the reactions in the model. Each parameter is of the form object.field. Any number of parameters can be specified, separated by spaces.", default = [] )
    parser.add_argument( '-s', '--stimulus', type = str, nargs = '+', metavar = 'args', action='append', help='Optional: Deliver stimulus as follows: --stimulus molecule conc time [conc time]... Each stimulus molecule may be followed by one or more [conc time] pairs. Any number of stimuli may be given, each indicated by --stimulus. Stimuli can overlap with the builtin stimuli, the values will apply from the time they are given till the builtin protocol delivers its own stimulus to override them.', default = [] )
    parser.add_argument( '-p', '--plot', action='store_true', help='Flag: when set, it plots the chem output, the original HillTau output, and the optimized HillTau output')
    parser.add_argument( "-t", "--tolerance", type = float, help = "Optional: tolerance for convergence of optimization.", default = 1.0e-6 )
    parser.add_argument( '-o', '--optfile', type = str, help='Optional: File name for saving optimized HillTau model. If not set, no file is saved.', default = "" )
    args = parser.parse_args()

    if len( args.builtin ) > 0:
        plotDt = min( plotDt, float( args.builtin[0][2] ) * stimRange[0] * 0.2 )
    stimVec = parseStims( args.stimulus, args.builtin, args.cyclic, args.dose_response )
    t0 = time.time()
    referenceOutputs = runMoose( args.chemModel, stimVec, args.monitor )
    t1 = time.time()
    print( "Completed reference run of '{}' in {:.2f}s".format( args.chemModel, t1 -t0 ) )

    mash = makeMash( args, stimVec, referenceOutputs )
    
    initParams = np.ones( len( mash.params ) )
    initRet = mash.doRun( initParams )
    bounds = [(0.01, 100.0)] * len( mash.params )

    x0 = initParams

    ret = minimize( mash.doEval, x0, method = "L-BFGS-B", tol = args.tolerance, bounds = bounds, callback = dotter )

    finalRet = mash.doRun( ret.x )
    print( "\n{:20s}   {}".format( "Object.field", "Scale factor" ) )
    for i, j in zip( mash.params, ret.x ):
        print( "{:20s}  {:4f}".format( i, j ) )

    print( "Timings: reference= {:.2f}s, optimization= {:.2f}s, HillTau Cumulative = {:.2f}s \nNumber of evaluations = {}, number of optimization iterations = {}, \nInitial score = {:3g}, Final score = {:3g}".format( t1 - t0, time.time() - t1, mash.simt, mash.numIter, ret.nit,  mash.doScore( initRet ), ret.fun ) )

    if len( args.optfile ) > 0:
        mash.dumpScaledFile( ret.x, args.optfile )


    if args.plot:
        for name, ref in referenceOutputs.items():
            hname = mash.molMap[ name ]
            fig = plt.figure( figsize = (6,6), facecolor='white' )
            ax = plotBoilerplate( xlabel = "Time (s)", title = hname )
            x = np.array( range( len( ref ) ) ) * plotDt
            ax.plot( x , 1000.0 * ref, label = "Mass action" )
            ax.plot( x , 1000.0 * np.array( initRet[hname] ), label = "HillTau" )
            ax.plot( x , 1000.0 * np.array( finalRet[hname] ), label = "HillTau opt" )
            ax.legend()
        plt.show()

if __name__ == '__main__':
    main()
```
