## Supplementary Code for "HillTau: A fast, compact abstraction for model reduction in biochemical signaling networks": README.txt

This directory contains Python scripts and models for generating most of the 
figures in the HillTau paper. 

README.txt			This file.

Figures.
File				Function			Depends on
fig1.py				make plots for fig 1		MOOSE, HillTau
fig1_supp.py			make supplementary figure 1	MOOSE, HillTau
fig2.py				make plots for fig 2		MOOSE, HillTau
fig3.py				make plots for fig 3		MOOSE, HillTau
fig5.py				make plots for fig 5		MOOSE, HillTau
fig5_supp1.py			make plots for supp fig 5.1	MOOSE, HillTau
fig5_supp2.py			make plots for supp fig 5.2	MOOSE, HillTau
fig5_supp3.py			make plots for supp fig 5.3	MOOSE, HillTau
fig5_supp4.py			make plots for supp fig 5.4:	HillTau, COPASI
				convert HillTau models to SBML	pycotools,
				and run them in COPASI,		pandas
				compare with HillTau originals
FIG5_SUPP4/			
fig6.py				Benchmark runs for fig 6	MOOSE, HillTau,
				Run repeatedly to get average	COPASI
				Should be run with C++ version
				of HillTau.
fig6_plotting.py		Plot fig6 using info from	-
				fig6.py
supp_fig7.py			Generate Supplementary fig7.	HillTau


Utility programs.
File				Function			Depends on
conv_kkit_to_sbml.py		Convert kkit models to SBML	MOOSE
insert_benchmark_events.py	Add events to SBML file, 	simplesbml
				used for some of the benchmarks

A few files from HILLTAU Github site are placed here for convenience.
Note that authoritative version is on GitHub.
Advisable to run with the compiled C++ version.
File				Function			Depends on
hillTau.py			HillTau Python version.		Python 3.x
hillTauSchema.json		Schema file for HillTau
ht2sbml.py			Convert HT models to SBML	HillTau, simplesbml
htgraph.py			Generate graphs of HT models	HillTau, graphviz
mash.py				Model Abstraction from SBML 	MOOSE, HillTau
				to HillTau

Directories
HT_MODELS/			HillTau models used in paper
KKIT_MODELS/			KKIT (Genesis format) models used in paper
SBML_MODELS/			SBML models used in paper.


			-----------------------------------
HillTau version alert: 
There is a _local_ version of hillTau.py in this directory to simplify 
running and installing. This local file should be renamed if you do a full 
HillTau install from the GitHub site, otherwise the local file will be the 
one that is called.
This will run slowly compared to the C++ HillTau version, and also it will 
not benefit from updates that will come to the GitHub site.

There is also a local version of mash.py. Again, the GitHub version will
be more current.

			-----------------------------------

Calls for running MASH optimizations
(You _really_ should use the compiled version of HillTau for this!)

Figure 2C
python mash.py KKIT_MODELS/fb_inhib.g HT_MODELS/fb_inhib_old.json -b input 0.001 80 -p -o fg_inhib.json -t 1e-9

Figure 2D
python mash.py KKIT_MODELS/kholodenko.g HT_MODELS/kholodenko_old.json -m MAPK_PP:output -s MAPK_PP/output 0 5000 -r MAPK.concInit MKKK.concInit -t 1e-9 -p -o kholodenko.json


Figure 3 CDE
python mash.py KKIT_MODELS/bcm.g HT_MODELS/bcm_old.json -m p_AMPAR:synAMPAR -b Ca 0.5e-3 20 -b Ca 5e-3 20 -p -t 1e-9 -o bcm.json

Figure 5 D-K
python mash.py KKIT_MODELS/acc92_fixed.g HT_MODELS/syn_prot_composite_old.json -a protein.tau protein.tau2 protein.KA protein.Kmod protein.Amod protein.gain protein.baseline -c Ca 10e-3 2 98 80 -s BDNF 3.7e-6 0 0.05e-6 8000 -d Ca 0.5e-3 2000 -d BDNF 1e-6 2000 -p -m protein -t 1e-9 -o syn_prot_composite.json

Fig 5 supp 1:
python mash.py KKIT_MODELS/acc92_fixed.g HT_MODELS/syn_prot_minimal_old.json -c Ca 10e-3 2 98 80 -s BDNF 3.7e-6 0 0.05e-6 8000 -d Ca 0.5e-3 2000 -d BDNF 1e-6 2000 -p -m protein -t 1e-9 -o syn_prot_minimal.json

Fig 5 supp 2:
python mash.py KKIT_MODELS/acc92_fixed.g HT_MODELS/syn_S6K_old.json -r BDNF.concInit Ca.concInit S6K.concInit TRKb.concInit aTRKb.tau2 -c BDNF 3.7e-6 1000 2000 10 -s Ca 2e-3 18000 0.08e-3 32000 -s BDNF 0.05e-6 32000 -d Ca 0.5e-3 2000 -d BDNF 1e-6 2000 -s Ca 0.08e-3 46100 -p -m S6K_thr_252:aS6K -t 1e-9 -o syn_S6K.json


Fig 5 supp 3:
python mash.py KKIT_MODELS/acc92_fixed.g HT_MODELS/syn_CaMKIII_old.json -a aCaMKIII.tau aCaMKIII.tau2 aCaMKIII.KA aCaMKIII.Kmod aCaMKIII.Amod aCaMKIII.baseline CaMKIII.concInit -c Ca 1e-3 20 100 5 -c Ca 10e-3 2 98 10 -s BDNF 3.7e-6 1400 0.05e-6 1800 -d Ca 0.5e-3 1000 -s Ca 1e-3 14000 -d BDNF 1e-6 1000 -p -m CaMKIII_CaM_Ca4:aCaMKIII -t 1e-9 -o syn_CaMKIII.json
