## Supplementary Code for "HillTau: A fast, compact abstraction for model reduction in biochemical signaling networks": supp_fig7.py

```
# 
# This program is free software; you can redistribute it and/or
# modify it under the terms of the GNU General Public License as
# published by the Free Software Foundation; either version 3, or
# (at your option) any later version.
# 
# This program is distributed in the hope that it will be useful,
# but WITHOUT ANY WARRANTY; without even the implied warranty of
# MERCHANTABILITY or FITNESS FOR A PARTICULAR PURPOSE.  See the GNU
# General Public License for more details.
# 
# You should have received a copy of the GNU General Public License
# along with this program; see the file COPYING.  If not, write to
# the Free Software Foundation, Inc., 51 Franklin Street, Fifth
# Floor, Boston, MA 02110-1301, USA.
# 

'''
*******************************************************************
 * File:            fig2.py
 * Description:
 * Author:          Upinder S. Bhalla
 *
 ********************************************************************/

/**********************************************************************
** This program uses HILLTAU to look at convergence of solutions with
** smaller timesteps.
**           copyright (C) 2021 Upinder S. Bhalla. and NCBS
**********************************************************************/
'''
from __future__ import print_function
import sys
import os
import json
import re
import argparse
import numpy as np
import matplotlib.pyplot as plt
import hillTau

t1 = 10
t2 = 50
t3 = 60
i1 = 1e-3

plotDt = 1

panelName = ["A", "B", "C", "D"]
color = ["C0", "C1", "C2", "C3", "C4", "C5", "C6"]

def plotBoilerplate( plotPos, reacn, xlabel = 'Time (s)', ylabel = 'Conc ($\mu$M)' ):
    panelTitle = panelName[plotPos-1]
    ax = plt.subplot( 4, 1, plotPos )
    ax.spines['top'].set_visible( False )
    ax.spines['right'].set_visible( False )
    '''
    ax.tick_params( direction = 'out' )
    '''
    ax.set_xlabel( xlabel, fontsize = 14 )
    ax.set_ylabel( ylabel, fontsize = 14 )
    ax.text( -0.32, 1, panelTitle, fontsize = 18, weight = 'bold', transform = ax.transAxes )
    #ax.text( 0.03, 1.03, reacn, fontsize = 12, transform = ax.transAxes )
    return ax

def runSim( dtList, ht, plotPos, stim = "input", output = "output", runtime = t3 ):
    ht = "HT_MODELS/" + ht
    ax = plotBoilerplate( plotPos, "fb_inhib", xlabel = "Time (s)" )
    jsonDict = hillTau.loadHillTau( ht )
    hillTau.scaleDict( jsonDict, hillTau.getQuantityScale( jsonDict ) )
    model = hillTau.parseModel( jsonDict )
    printMinTau( model, ht )
    ovec = []
    for idx, dt in enumerate( dtList ): 
        model.dt = dt
        model.reinit()
        model.internalDt = dt # We override internalDt to do accuracy check
        #print( "Timesteps = ", dt, model.internalDt )
        inputMolIndex = model.molInfo.get( stim ).index
        outputMolIndex = model.molInfo.get( output ).index

        model.advance( t1 )
        model.conc[inputMolIndex] = i1
        if ( runtime < t3 ):
            model.advance( runtime - t1 )
        else:
            model.advance( t2 - t1 )
            model.conc[inputMolIndex] = 0
            model.advance( t3 - t2 )
        plotvec = np.transpose( np.array( model.plotvec ) )
        x = np.array( range( plotvec.shape[1] ) ) * dt
        reacn = "this is ht"
        htvec = np.array( plotvec[outputMolIndex] )
        ax.plot( x , 1000 * htvec, label = "dt= " + str(dt), color = color[idx], linestyle = ":" )
        #print( len( htvec ), round(dtList[-1]/dt) )
        ovec.append(htvec[::round(dtList[-1]/dt) ] )
        ax.plot( x[::round(dtList[-1]/dt)] , 1000 * ovec[-1], marker = ".", color = color[idx], linestyle = "none" )

    ax.legend()
    for v, dt in zip(ovec, dtList):
        ml = min( len( v ), len( ovec[0] ) )
        #print( "ML = ", ml, v )
        dy = v[:ml] - ovec[0][:ml]
        print( "{} @ dt= {}: normalized rms diff ={:.4f}".format( ht, dt, np.sqrt( np.mean( dy * dy )) / np.max( ovec[0] )) )
        


def runOsc( dtList, ht, plotPos ):
    ht = "HT_MODELS/" + ht
    runtime = 3000
    ax = plotBoilerplate( plotPos, "Oscillator", xlabel = "Time (s)" )
    jsonDict = hillTau.loadHillTau( ht )
    hillTau.scaleDict( jsonDict, hillTau.getQuantityScale( jsonDict ) )
    model = hillTau.parseModel( jsonDict )
    printMinTau( model, ht )

    ovec = []
    for idx, dt in enumerate( dtList ): 
        model.dt = dt
        model.reinit()
        model.internalDt = dt
        #print( "Timesteps = ", dt, model.internalDt )
        outputMolIndex = model.molInfo.get( "output" ).index

        model.advance( runtime )
        plotvec = np.transpose( np.array( model.plotvec ) )
        x = np.array( range( plotvec.shape[1] ) ) * dt
        reacn = "this is ht"
        #ax = plotBoilerplate( "H", plotPos+4, reacn, xlabel = "Time (s)" )
        #ax.set_ylim( 0, 0.4 )
        htvec = np.array( plotvec[outputMolIndex] )
        ax.plot( x , 1000 * htvec, label = "dt= " + str(dt), color = color[idx], linestyle = ":" )
        #print( len( htvec ), round(dtList[-1]/dt) )
        ovec.append(htvec[::round(dtList[-1]/dt) ] )
        ax.plot( x[::round(dtList[-1]/dt)] , 1000 * ovec[-1], marker = ".", color = color[idx], linestyle = "none" )

    ax.legend()
    for v, dt in zip(ovec, dtList):
        dy = v - ovec[0]
        print( "{} @ dt= {}: normalized rms diff ={:.4f}".format( ht, dt, np.sqrt( np.mean( dy * dy )) / np.max( ovec[0] )) )

def printMinTau( model, name ):
    tau = 1e6
    for r in model.reacInfo.values():
        tau = min( tau, r.tau, r.tau2 )
    print( "Model {}.   Minimum tau = {}".format( name, tau ) )
    

def main():
    fig = plt.figure( figsize = (6,12), facecolor='white' )
    fig.subplots_adjust( left = 0.18 )
    runSim( [0.1, 0.2, 1.0, 2.5, 5.0], "fb_inhib.json", 1 )
    runSim( [0.1, 0.2, 1.0, 2.5, 5.0], "ff_inhib.json", 2 )
    runSim( [0.1, 0.2, 1.0, 2 ], "bcm.json", 3, stim = "Ca", output = "synAMPAR", runtime = 20 )
    runOsc( [1, 6, 10, 60, 300],"kholodenko.json", 4 )

    plt.tight_layout()
    plt.show()


if __name__ == '__main__':
    main()
```
